# Boundary effects cause false signals of range expansions in population genomic data

**DOI:** 10.1101/2023.12.06.570483

**Authors:** Petri Kemppainen, Rhiannon Schembri, Paolo Momigliano

## Abstract

Studying range expansions (REs) is central for understanding genetic variation through space and time as well as for identifying refugia and biological invasions. Range expansions are characterized by serial founder events causing clines of decreasing diversity away from the center of origin and asymmetries in the two-dimensional allele frequency spectra. These asymmetries, summarized by the directionality index (ψ), are sensitive to REs and persist for longer than clines in genetic diversity. In continuous and finite meta-populations, genetic drift tends to be stronger at the edges of the species distribution. Such boundary effects (BEs) are expected to affect geographic patterns in ψ as well as genetic diversity. With simulations we show that BEs consistently cause high false positive rates in equilibrium meta-populations when testing for REs. In the simulations, the absolute value of ψ (|ψ|) in equilibrium data sets was proportional to the fixation index (*F_ST_*). By fitting signatures of REs as a function of ɛ=|ψ|/*F_ST_* and geographic clines in ψ, strong evidence for REs could be detected in data from a recent rapid invasion of the cane toad, *Rhinella marina*, in Australia, but not in 28 previously published empirical data sets from Australian scincid lizards or the Indo-Australasian blacktip shark that were significant for the standard RE tests. Thus, while clinal variation in ψ is still the most sensitive statistic to REs, in order to detect true signatures of REs in natural populations, its magnitude needs to be considered in relation to the overall levels of genetic structuring in the data.

## Introduction

Species ranges are seldom static through time. For instance, when new suitable habitat becomes available a species may colonize previously unpopulated areas through range expansions (RE). Studying the expansion of populations across a landscape is not only central for understanding the demographic histories of natural populations (including humans), but is also necessary for understanding and predicting disease outbreaks, biological invasions and the spread of a native species across novel geographic regions made newly suitable by climate change (O’Reilly-Nugent et al. 2016; Ogden et al. 2019; Poland et al. 2021; Alves et al. 2022; Selechnik, Richardson, Shine, DeVore, et al. 2019; Ioannidis et al. 2021; Zhan et al. 2014; Zhang et al. 2022; Finch et al. 2021). Range expansions imply sequential founder events and leave transient signatures on the distribution of genetic diversity across the meta-population including clines of decreasing genetic diversity (e.g. expected heterozygosity, *H_E_*) away from the center of origin (Ramachandran et al. 2005) and asymmetries in the two-dimensional site frequency spectra (2D- SFS) between populations (Peter and Slatkin 2015, 2013).

Due to the increased genetic drift associated with founder events, newly colonized geographic locations are expected to have an excess of intermediate frequency allelic variants compared to the population they originated from. This can be estimated as the directionality index, ψ, defined for pairs of populations (*P_1_* and *P_2_*) as the number of SNP fixed for the derived allele in *P_1_* and heterozygous in *P_2_* minus the number of SNP fixed for the derived allele in *P_2_* and heterozygous in *P_1_* divided by the total number of sites segregating in both populations. Previous simulation work has shown clines in ψ to be more sensitive and robust to detecting signatures of REs and estimating their origins compared to methods based on clines in genetic diversity (Peter and Slatkin 2015, 2013), where the expansion origin is expected to be the location with the strongest positive correlation between geographic distance and ψ. This approach has been increasingly applied in recent population genetic studies as it only requires one diploid individual to be sampled per population (Zhan et al. 2014; Maisano Delser et al. 2019; Prior et al. 2020; Fifer et al. 2022; Hemstrom et al. 2022; Lesturgie et al. 2023; Singhal, Wrath, and Rabosky 2022; Walsh et al. 2022; Ioannidis et al. 2021; He, Prado, and Knowles 2017; Jaya et al. 2022). A further potential benefit of the methodology introduced in Peter and Slatkin (2013, 2015) is the use of the Time Difference of Arrival (TDoA) - a ranging technique regularly used in the Global Positioning System (GPS) that allows for the inference of range expansion origins also from unsampled geographic regions. A more advanced method to estimate the origins of REs based on Approximate Bayesian Computation (ABC) is also now available (He, Prado, and Knowles 2017)

Range expansions, however, are not the only process to produce asymmetries in the 2D-SFS. Differences in effective population sizes (*N_e_*’s) and/or asymmetric migration can produce similar patterns (Gutenkunst et al. 2009; Marchi and Excoffier 2020). More importantly, because population centers on average receive a more genetically diverse set of migrants (from all directions of the distribution range), range margins typically exhibit lower genetic diversity, resulting in clines of decreasing diversity away from the center of a species range (Eckert, Samis, and Lougheed 2008; Wilkins and Wakeley 2002). Since such boundary effects (BEs) are ultimately driven by increased levels of genetic drift (smaller *N_e_*’s) at the edges, BEs will cause asymmetries in 2D-SFS as well (Gutenkunst et al. 2009). Despite the fact that clear signatures of BEs were observed in the geographic patterns of genetic diversity in simulated equilibrium isolation-by-distance populations in Peter and Slatkin (2013, 2015; and to some extent also in spatial patterns in ψ), no excessive false positive rates for statistical tests for REs were reported in these studies. However, the original simulations in Peter and Slatkin (2013, 2015) were based on simple Wright-Fisher models and employed a limited range of parameter values. Since the power of the tests used for rejecting the null-hypothesis of ψ≠0 depends on the number of segregating sites and only 1000 independent SNPs were used (Peter and Slatkin 2013), the lack of elevated false positives rates despite signs of BEs may also have been a matter of statistical power. Furthermore, the TDoA method relies on testing for correlations between two matrices (geographic distance and ψ) not accounting for the non-independence among the pairwise measures raising the possibility of severe *p*-value inflation (Peter and Slatkin 2015). Yet once the null-hypothesis of ψ≠0 has been rejected, finding significant correlations in TDoA have regularly been used to further support REs in population genomic data (Maisano Delser et al. 2019; Jaya et al. 2022; Peter and Slatkin 2015, 2013; Singhal, Wrath, and Rabosky 2022).

In this study, we tested the extent to which BEs under equilibrium conditions result in similar clinal variation in ψ and genetic diversity as expected during REs. We used spatially explicit, individual based non-Wright-Fisher 1D and 2D stepping stone simulations as well as simulations in two-dimensional continuous space with age structure and overlapping generations in heterogeneous seascapes modeled after the grey reef sharks in the coral-triangle (Robbins et al. 2006; Boussarie et al. 2022). We further demonstrate that the mean |ψ|-values in equilibrium populations is proportional to the overall levels of genetic structuring in the data as measured by the fixation index (*F_ST_*) and test to what extent the levels of ψ, *F_ST_*and clinal variation ψ independently and jointly can predict genetic signatures of REs in the simulated data using linear models. These models were then tested on 28 empirical data sets of Australian scincid lizards with large numbers of significant ψ-values and strong geographic clines in ψ but with no known historical records of range expansions (Singhal et al. 2022), on a previously analysed blacktip reef shark data from the Indian and Pacific Oceans (Maisano Delser et al. 2019), as well as on data from the recent and rapid invasion of the cane toad in Australia (Selechnik et al. 2019) that functioned as a positive control.

## Materials and methods

### 1D *and 2D simulations*

For the first set of simulations we aimed to add biological realism to the 1D and 2D stepping stone simulations presented in Peter and Slatkin (2013, 2015) by using individual based non-Wright fisher models implemented in SLiM 4 (Haller and Messer 2023). Based on recipe 16.19 in the SLiM manual (v. 4.0.1), we modeled a sexually reproducing hermaphroditic species with no selfing and non-overlapping generations, and with population size determined by negative frequency dependence (simulation parameters are summarized in Table 1). The models consisted of N=81 demes (d_1_,d_2_,.. d_81_) connected either as a single chain (1D) or arranged in a 9×9 matrix of demes (2D), with each deme connected by migration with adjacent demes (Fig. 1 a,b). The number of migrants between adjacent demes were drawn from a binomial distribution with a probability M for each individual in the source deme, with three levels resulting in low, medium or high gene flow between demes (Table 1). Since each deme in the 1D models can be connected to a maximum of two other demes, whereas in the 2D models a deme can be connected to a maximum of four demes (Fig. 1 a,b), the levels of M were chosen so that patterns of genetic isolation-by-distance (IBD) in 1D and 2D models would be more comparable (Table 1). Initially a single deme was allowed to reach mutation drift equilibrium either at the beginning or the middle of the stepping stone chain (1D) or in one of the corners or the middle of the 2D matrix (d_1_), after which the remaining (previously empty) populations were allowed to be colonized. The number of offspring per individual was drawn from a Poisson distribution with mean=1.04 such that after initial colonization the population size in a given deme would increase until limited by negative density dependence to its carrying capacity of K=1000. Thus, the speed of the RE was proportional to M. A single chromosome was simulated with a size L and mutation rate μ adjusted (Table 1) such that a minimum of 50k polymorphic SNPs would be available for analyses after REs were completed (at t_0_), when the total number of individuals in the meta- population reached 98% of K*N. Recombination rate r was uniform across the chromosome with r=μ which allowed linkage disequilibrium (LD) to decline rapidly with distance along the chromosome. After t_0_, genotypic data were saved for five individuals from each deme for downstream analyses at 11 time points ranging from 100 to 128k simulation cycles (equal to generations and years) post t_0_. While 128k generations was not sufficient for genetic diversity to reach equilibrium (overall genetic diversity did not reach a plateau at the end of the simulations) it was nevertheless sufficient to eliminate all signals of the REs (i.e. *F*_ST_ and *ψ* reached equilibrium). Thus, here we consider the population at the end of each simulation as the null model for REs, where the balance between mutation, drift and gene-flow outweighs the effects on patterns of genetic variation relative to the RE. As a contrast, we further included a fully panmictic model where all 81 demes had an equal probability of M=0.25 to exchange migrants with any other deme in the meta-population. All segregating SNPs were initially saved for downstream analyses but only 50k SNPs were subsequently sampled for the final data set (minor allele frequency, *maf*=1/2n where n is the number of individuals). Ten replicate simulations were run for each parameter combination of origin O (edge or center) and M (Table 1) for both 1D and 2D models as well as for the single parameter combination for the panmictic model.

**Figure 1.**
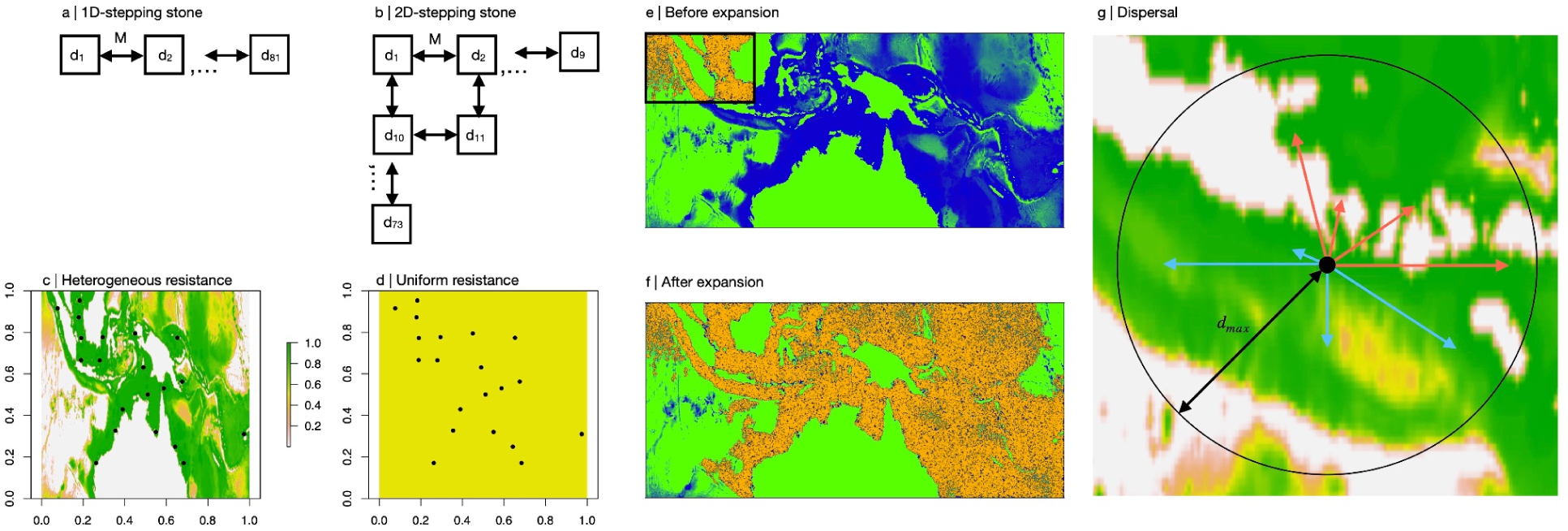
Simulation overview. The layout of demes in the 1D and 2D stepping stone simulations are shown in (a) and (b), respectively. The heterogeneous (HL) and uniform (UL) landscapes for the continuous space simulations for the coral triangle are shown in (c) and (d) with the population coordinates indicated. In (e) and (f) the distribution of individuals in HL are shown for an equilibrium population before range expansion (boundaries shown by the black box) and after range expansion, respectively. To model dispersal in the continuous space simulations, eight possible new positions for each individual were drawn a maximum of *d_max_* away from the old position (at the center of the circle). Any paths crossing land and any new position outside the map (c) were disregarded (red paths) and among the remaining possible new positions (blue), one was selected as the new position using the inverse of total resistance across the path as weight.

**Table 1.** Simulation parameters.

|  |  |  |
| --- | --- | --- |
| 1D and 2D stepping-stone |  |  |
| N | Number of demes | 81 |
| K | Carrying capacity per deme | 1000 |
| D | Number of dimensions | 1; 2 |
| M | Migration probability | ID: 0.01; 0.05; 0.1<br>2D: 0.002; 0.005; 0.01 |
| O | Origin of expansion | 1; 41 |
| $\mu$ | Mutation rate | 1.5e-7 |
| L | Chromosome length (bps)<br>(Uniform recombination) | 1e6 |
| r | Recombination rate | 1.5e-7 |
| 2D continuous-space |  |  |
| $d_{\max}$ | Maximum dispersal distance | 20; 10; 5 |
| R | Resistance map | Heterogeneous (HL); Uniform (UL) |
| X | Type of expansion | RE (range expansion);<br>DE (demographic expansion) |
| $\mu$ | Mutation rate | 1e-8 |
| L | Chromosome length (bps)<br>(recombination map:<br>*D. melanogaster* ChrII) | 2.3e6 |
| r | Recombination rate | 1e-8 |

### 2D *continuous-space simulations*

Based on ψ and geographic clines of genetic diversity, several recent papers on blacktip (*Carcharhinus limbatus*) and grey reef sharks (*Carcharhinus amblyrhynchos*) throughout the Indian and Pacific oceans, have inferred that there is sufficient evidence for REs in these two species (Maisano Delser et al. 2019; Lesturgie et al. 2023; Walsh et al. 2022; Boussarie et al. 2022) in support for the hypothesis that the Indo-Australasian archipelago is a center of origin for marine biodiversity. To test to what extent boundary effects may cause similar geographic patterns of genetic diversity, we loosely modeled 2D continuous-space simulations of the grey reef sharks in the coral triangle based on recipes 15.11 (biogeographic landscape model), 16.10 (spatial competition and mate choice), 16.2 (age structure) and 6.1.2 (heterogeneous recombination rates) in the SLiM manual. This allowed us to simulate a dioecious and iteroparous organism with overlapping generations in a heterogeneous seascape (with realistic recombination rate variation across a chromosome), where both the fitness and dispersal distance was proportional to the habitat quality in which the individual resides in.

Because the TDoA approach used to estimate the origins of range expansions assumes equal habitat suitability across space and time it is important to assess how isolation-by-resistance (IBR) models that incorporate the effects of heterogeneous habitats on gene flow (McRae and Beier 2007) may affect the accuracy of this method. Boussarie et al. (2022) showed that both bathymetry (sea depth) and distance from the closest coral reef best explained the patterns of genetic connectivity among populations of grey reef sharks in the coral triangle (samples collected across the Indian and Pacific oceans). The heterogeneous fitness landscape in our 2D continuous-space model was therefore based on the resistance map (398 by 855 matrix) from Boussarie et al. (2022) that produced the best IBR fit in the grey reef shark data (Fig. 1c) where the value of each grid point (physical position on the map) gives the habitat quality q=[1,0], where 0 represents land or depths <4km and 1 represents coral reef. With age structure, negative density dependence and spatial competition the individual fitness is given by:

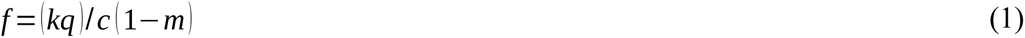

where m=[0.7, 0.0, 0.0, 0.25, 0.5, 0.75, 1.0] is the normalized age specific mortality, for ages 0-6 (reproduction starting at age one), c is the rescaled strength of competition felt by the individual (see recipe 16.10 for details) and k is a scaling parameter to control the total number of individuals in the simulations in order to keep run times manageable. When the population is at equilibrium i.e. when census population size (*N_c_*) is approximately equal to K, c will also be approximately equal to k. In all simulations k=100 which resulted in K∼7e^5^. Thus, the highest fitness could be reached for 1-2 year old (1-m=1) individuals residing in coral reefs (q=1), at an expansion front where population density was still low (c<k). The number of offspring was drawn from a Poisson distribution with mean 1, but since the organism was iteroparous, this ensured population growth whenever *N_c_*<K, for instance at the edges of an expansion, thus simulating a large species with slow reproduction.

The dispersal of individuals was modeled as follows. First, the direction of the dispersal was determined by randomly drawing eight directions. Second, for all directions the dispersal distance *d* was drawn from a standard uniform distribution multiplied by *q* and *d_max_*, the maximum possible distance an individual can disperse each simulation cycle (here equivalent to years but not generations), resulting in eight potential new positions to choose from. Third, the total resistance (*s*) for each new position was calculated as the sum of all grid points along the straight path from the old position to the new positions (the maximum resistance allowed was *s_max_*=8). If any of the grid points crossed by this line included land or deep sea (*q*=1) the new position was disregarded, likewise when the new position was outside the boundaries of the map (indicated by red in Fig. 1g). This kept individuals from crossing land and avoided severe BE’s around the map borders and borders between sea and land (or deep sea). From the remaining potential new positions for an individual (indicated in blue in Fig. 1g), one was selected using *w*=1-*s*/*s_max_* as a weight, i.e. the most likely new position was the one with the lowest total resistance on the path from the old position to the new position. If, among the eight possible new positions none were viable options (all paths crossing land or deep sea, or s>*s_max_*), the fitness of the individual was set to zero in effect killing the individual. This equates to absorbing boundaries, which ensures that population density remains proportional to habitat quality in the simulations across the whole map (with reprising/reflective boundaries, population density at edges would have been higher, which is not desirable) and is expected to result in BEs (Wilkins and Wakeley, 2002). The strength of IBR was ultimately determined by the parameter *d_max_* with three levels resulting in low, medium and high gene flow (Table 1), that also determined the speed of expansions, i.e. lower gene flow resulting in slower expansions.

As a contrast to the IBR model described above, we also simulated data using a map where resistance was uniform (UL) i.e., a standard IBD model. The census population in the simulations was a function of the mean resistance of all grid points on the map. Therefore, using the mean of all grid points (0.516) from the HL map in UL ensured that K would be similar, with the difference that population density in HL was highly patchy and centered around coral reefs and corridors of shallow areas along coastlines, whereas in UL the average local population density was uniform across the map. The simulations began with an equilibrium population (generated by using 20k cycle burn-ins) with approximately 1/10th of the map (*K*∼7e^4^, Fig. 1e) being populated before allowing for individuals to expand across the remainder of the map. As a contrast to the RE model, we also allowed the individuals from the burn-in to immediately seed the entire map simulating a demographic expansion (DE) without a spatial component.

All polymorphic SNPs from 2000 randomly chosen individuals were saved at 600, 800, 1000, 2500, 5000, 10000, 20000, 40000 and 80000 cycles after expansions started for downstream analyses. This was sufficient for RE and DE models to converge to similar estimates (as simulations proceeded) for all population genetic summary statistics except genetic diversity. Therefore, the data set sampled at the end of the simulations were appropriate as a null model for testing REs. Based on general patterns of genetic diversity and population densities, 20 population cores (coordinates) were selected (Fig. 1 b) and for each of them the 10 closest individuals were chosen as the population sample for the final data set (n=200).

In order to add some level of realism to the recombination landscape, in the absence of accurate recombination maps for shark genomes, the recombination map followed that of *Drosophila melanogaster* ChrII (Comeron, Ratnappan, and Bailin 2012) and μ was adjusted such that a minimum of 50k polymorphic SNPs could be sampled for all data sets, also here with r=μ (Table 1). Ten replicate simulations were run for each parameter combination of *d_max_*, type of expansion (RE or DE) and habitat heterogeneity (HL or UL; Table 1).

### Testing range expansions and finding origins

Tests for range expansions and estimation of the center of origin followed the same approach for the stepping stone models and the continuous space models unless otherwise stated. As in recent studies (Singhal, Wrath, and Rabosky 2022; Jaya et al. 2022; He, Prado, and Knowles 2017), we used the original functions and pipelines from the R-package rangeExpansion (v.0.0.0.9000; Peter and Slatkin 2013, 2015; https://github.com/BenjaminPeter/rangeexpansion) to estimate ψ and the centers of origin. Some modifications of the original code were, however, necessary to correct some bugs, streamline the pipelines and improve computational speed. Most importantly, the polarity of the ψ-matrix produced by the original code is reversed, resulting in the most likely origin to be estimated where the genetic diversity is the lowest (i.e. the most recently colonized population), instead of the highest (Supplementary File 1). From the simulation output we prepared a genotype and a coordinate file and used the function *preparedload.data.snapp* followed by *make.pop* to prepare the raw data for range expansion analyses. To check independence of SNPs, LD between all pairs of adjacent SNPs was estimated using function *snpgdsLDMat* from the R-package SNPrelate as the squared correlation coefficient *r^2^*(Zheng et al. 2012). While in Peter and Slatkin (2013) a block jackknifing approach was used to account for the non-independence of loci when estimating the significance of ψ, R-package rangeExpansion does not include any function to perform this operation. However, since only ∼0.5% (stepping stone models) or ∼5% (continuous space models) of all adjacent SNPs along the chromosome showed *r^2^*>0.2 in of the simulated data sets, SNPs could to a large extent be considered as independent. Therefore, we tested significance of ψ by first projecting down the 2D-SFS to one diploid individual per population and testing whether absolute frequencies of alleles polymorphic in both populations deviated from 0.5 using a binomial test (Peter and Slatkin 2013). We used the custom function *get.all.psi.mc.bin* (a modification of *get.all.psi* from rangeExpansion), that performed the binomial test but also improved the speed of the ψ estimations by a minimum of one order of magnitude.

Next, the function *prep.tdoa.data* was used to prepare data for the TDoA estimations of range expansion origins using the function *single.origin* (Peter and Slatkin 2015) as described in Supplementary File 2. In short, the map was divided into a 100×100 grid of evenly spaced coordinates and a linear regression was used to find the grid point that shows the strongest positive correlation between ψ and the difference in the geographic distance from this grid point to each of the two populations (for which the ψ was estimated). As an alternative to ψ, we used the same approach using the difference in *H_E_* between pairwise populations (Δ*_het_*) where the correlation is instead expected to be negative (Ramachandran et al. 2005). Note that TDoA, as it is implemented in the rangeExpansion package is not equivalent to the original implementation given in Peter and Slatkin (2013) and relies on correlations between two pairwise distance matrices (geographic distance and ψ) without correcting for non-independence among the data points and is thus expected to lead to high false positive rates (by overestimating the degrees of freedom; Supplementary File 2). Therefore, we also used the more traditional and conservative (non-TDoA) approach where a linear regression was used for the correlation between the pairwise distances from a focal population and all other populations in the data set and the corresponding ψ (or negative correlation with Δ*_het_*; Ramachandran et al. 2005) where the degrees of freedom are not overestimated. In the stepping-stone models (but not for the continuous space models), the grid points for the TDoA analyses corresponded to the coordinates for the populations so the estimated origin always overlapped with a sampled population for both the TDoA and the non-TDoA approaches. As in Peter and Slatkin (2013) we used the root mean squared error (RMSE) of the Euclidean distance between the estimated and the true origins to determine the accuracy of the TDoA and the non-TDoA methods. Since the 2D models were symmetric with range expansion starting from one of the corners (or from the middle), we used the mean across the x and y coordinates of the population matrix for comparing spatial patterns of ψ and *H_E_*with the 1D models. In the continuous space simulations, the center of the geographic regions populated during the burn-ins was used as the true range expansion origin for the RMSE estimates.

### Predicting boundary effects at equilibrium in simulated data

Since BEs in equilibrium populations are caused by increased genetic drift (declining *N_e_*’s) towards the edges of finite populations (Wilkins and Wakeley 2002), we expect them to affect geographic patterns of ψ across the meta-population as well (Gutenkunst et al. 2009; Peter and Slatkin 2015, 2013). The strength of the correlation between geographic distance from the population center and genetic diversity due to BEs in equilibrium populations is proportional to the genetic structuring in the data (Wilkins and Wakeley 2002). It is thus reasonable to expect a similar relationship between geographic clines in ψ and population connectivity as well. If this is true, rescaling ψ by *F_ST_* is expected to normalise the levels of ψ in equilibrium meta-populations across different levels of genetic structuring. We thus define the scaled ψ as 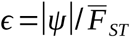, where 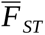 (hereafter simply *F_ST_*) is the mean pairwise *F_ST_* (Weir and Cockerham 1984) as estimated by function *snpgdsFst* from R-package SNPRelate (Zheng et al. 2012). Because when *F _ST_ →*0 *, ɛ→∞*, all *F_ST_* values were censored to a minimum value of *F_ST_*=0.001 across the panmictic simulations (*F_ST_* was above 0.001 in all other simulations). The expected *ɛ* for a population at equilibrium is here defined as *ɛ _eq_* and while *ɛ _eq_* cannot be known in empirical data, it can be estimated from the data sets sampled at the end of our simulations where |ψ| and *F_ST_* (although not necessarily genetic diversity) are expected to be at (or close to) equilibrium. Thus, the *ɛ _eq_* estimator can here be considered as the null-distribution and we are interested in testing the hypothesis that *ɛ* > *ɛ_eq_*in a given data set. Assuming that no other evolutionary phenomena except BEs cause elevated levels of ψ at equilibrium across the meta-population, we define the effect size for the genetic signature of a RE as *Ε*=*ɛ* / *ɛ _eq_* where *Ε*>1 indicates that the observed*ɛ* is higher than what can be expected at equilibrium due to BEs. Based on the simulated data, where the true *ɛ* _eq_ can be estimated with some confidence, we can thus model *E* as:

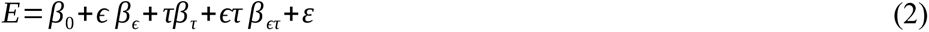

where*τ* is the effect size (strongest positive *r^2^*between any population pair in the data) from TDoA or the non-TDoA methods, *β*_0_ is the intercept, *β_ɛ_*and *β_τ_*are the regression coefficients for *ɛ* and *τ*, respectively, *ɛτ β_ɛτ_* represents the interaction between these two variables and *ε* is the residual variance. Due to heteroscedasticity in *ɛ*, the estimation of standard errors were weighted by 1/*ɛ*. Since only *ɛ* and *τ* can be estimated from empirical data, the primary focus of the model was to determine how well *E* can be predicted across multiple simulation models, migration/dispersal parameters and type of expansion (a function of how consistent *ɛ _eq_* is across the different simulation models). To test to what extent there is a significant genetic signature of REs in the data - beyond what can be expected by BEs at a given level of genetic structuring - we can find the lower boundaries for the prediction intervals satisfying *E*>1 from this model, for given confidence levels, α.

### Re-analyses of 30 empirical data sets

We re-analysed reduced representation sequencing data from 8-73 individuals (mean=16.5) and 3821-87570 SNPs from 28 empirical scincid lizards data sets with various distribution ranges across the Australian continent, where previous studies have found evidence of REs based on ψ and clines thereof (Singhal et al. 2022). Second, we re-analysed a population genomic data from the blacktip shark sampled across the Indian and Pacific from Masano Delser et al. (2019) - a study supporting the hypothesis of the Indo-Australasian archipelago being a center of origin for marine biodiversity (431257 SNPs from 144 individuals and 13 populations). Lastly, we analysed population genomic data from a recent rapid range expansion with a known origin from historical data: the invasion of the cane toad, *Rhinella marina*, in Australia. Historical records document the introduction of the cane toad to Gordonvale, North Queensland, in 1935 (Sabath, Boughton, and Easteal 1981; Easteal 1981), and they have since spread rapidly across the northern half of the continent to become a widespread and destructive pest species (Phillips and Shine 2004). Raw reads from RNAseq data from cane toad brains sampled across northern Australia (Selechnik, Richardson, Shine, Brown, et al. 2019) were accessed from the NCBI Short Read Archive PRJNA479937 and trimmed for quality and adapter contamination using Trimmomatic v.0.32 (Bolger, Lohse, and Usadel 2014). Reads were then mapped to the reference transcriptome of the closely related species *Rhinella arenarum*, the Argentine toad (Ceschin et al. 2020) with bwa mem (Li and Durbin 2009). SNPs were called with bcftools mpileup (Danecek et al. 2021) and polarised against the outgroup reference transcriptome to obtain the ancestral state. SNPs were filtered for quality and individuals with less than 90% call rate were excluded, after which one SNP per reference transcript was randomly chosen to minimise LD between SNPs. Following filtering, the dataset included 58 individuals and 18658 SNPs. All summary statistics for the empirical data were then estimated as for the simulated data.

## Results

### 1D and 2D stepping stone models

The mean *F_ST_* for the two most distant populations in the 1D simulations were 0.17, 0.29 and 0.63 for high, medium and low levels of gene flow, respectively and the corresponding values for the 2D simulations were 0.13, 0.21, and 0.42. The simulations confirmed that ψ is more sensitive and retains the signal of a RE longer than *H_E_* (Fig. 2), but, in contrast to Peter and Slatkin (2013, 2015), we also found clear patterns consistent with BEs not only for *H_E_,* but also for ψ (Fig. 2). The population with the highest genetic diversity and lowest ψ at equilibrium (>10k cycles post t_0_) was always found at the center of the meta-population, even when the expansion started from the edge (Fig. 2b). Furthermore, when the expansion started from the edge, the lowest ψ and highest *H_E_* were never found where the expansion started (*d_1_*), but was instead increasingly biased towards the center with increasing times in a “wave like” manner (Fig. 2). Furthermore, the stronger the range expansion signal was in the beginning (t_0_+100 cycles), the stronger the boundary effect was at the end of the simulations (t_0_+128k cycles; measured as the maximum difference in ψ or *H_E_* between any two populations; 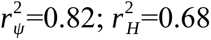; Fig. 2).

**Figure 2.**
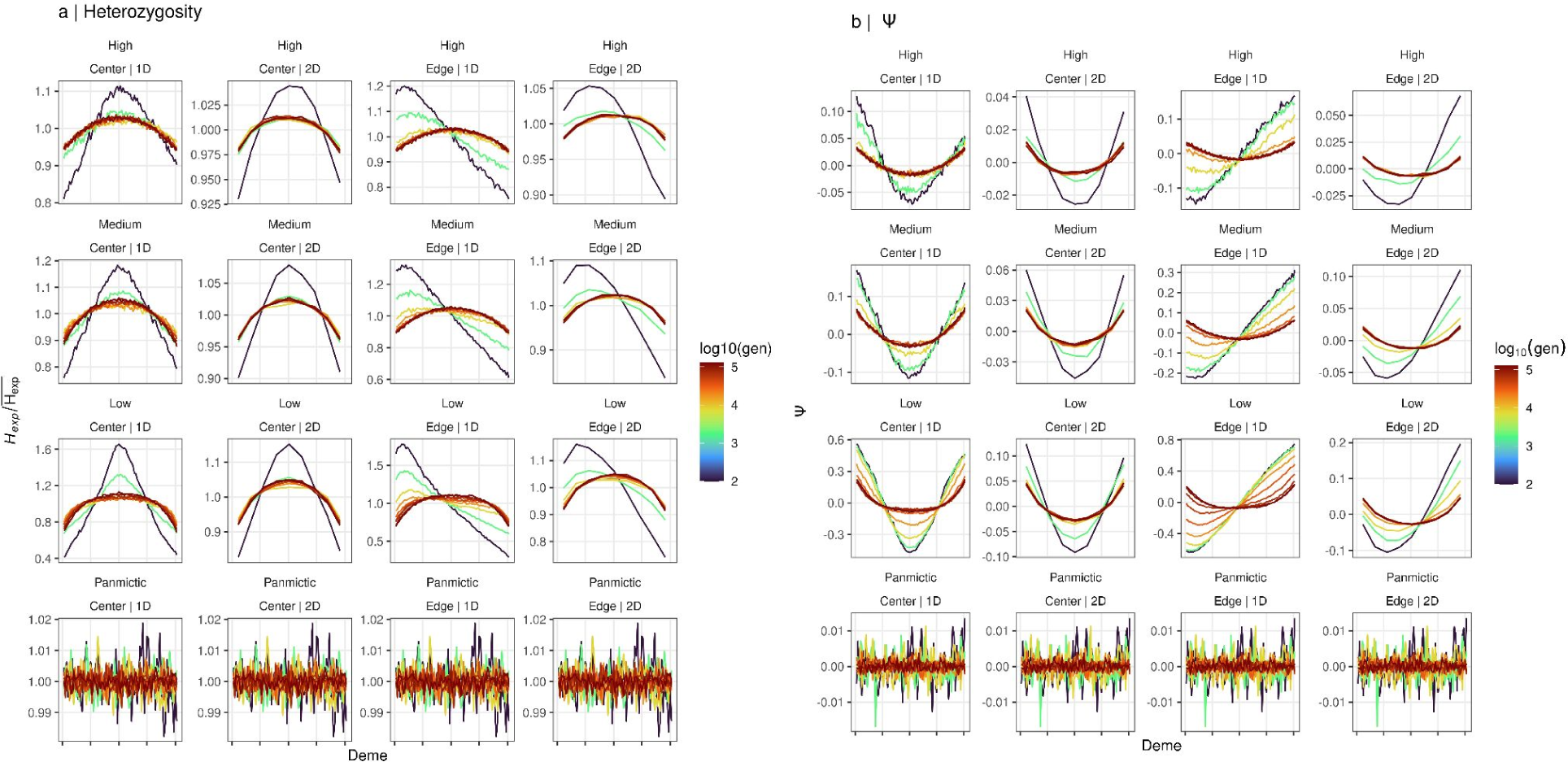
Spatial variation in ψ and genetic diversity in 1D and 2D stepping-stone simulations. Shows (a) mean standardized *H_E_* (n=10) and (b) ψ for demes *d_1_*to *d_81_* for 1D models and the mean of the x and y dimensions for the symmetric 2D models. Color represents the number of cycles (generations) since expansion was completed on a *log_10_* scale. Simulation parameters are indicated above each panel (low, medium and high gene flow with expansions starting from the center or from the edge, for 1D and 2D simulations, respectively). Results for panmictic simulation are shown for reference (here dimensionality and origin of expansion are irrelevant). Note the different y-scales for each row of panels.

Using 50k SNPs, the binomial test for ψ was always significant for >5% of comparisons (subsequently also resulting in non-zero significant ψ-values after Bonferroni correction for multiple testing), with the exception of the scenarios with high gene flow, and only for the short time interval where the signature of the REs diminished but the signature of the BEs was not yet strong (Supplementary Fig. 1). No ψ-values were significant under the panmictic scenarios. After an initial reduction (approximately after 10-40k cycles post t_0_), the proportion of rejected null-hypotheses increased (Supplementary Fig. S1a). This was not a function of increasing levels of asymmetries in the data (Supplementary Fig. S1b), but instead a function of increased statistical power due to larger numbers of SNPs segregating in both populations available for the binomial test (Supplementary Fig. S1c). This pattern was true also for medium and low levels of gene flow, such that the proportion of significant ψ values was a poor predictor of whether a signal of RE still remained in the data or if the signal was solely caused by a BE.

Once the null hypothesis of ψ≠0 has been rejected, to find support for REs, clinal variation in ψ should also be demonstrated (Peter and Slatkin 2013, 2015). However, it is important to note that significant correlations between geographic distance and ψ will almost always be found with the TDoA approach (Peter and Slatkin 2013). Indeed, close to 100% of all TDoA regressions were significant for all data sets, except when the meta-population was panmictic, in which case on average ∼50% of all tests were significant (Supplementary Fig. S2a). In the panmictic simulations the strongest positive correlation between geographic distance and ψ was significant in 82% of the data sets at α=0.05. When using α=1e-07, this number was still >50%. Except when populations were panmictic, the non-TDoA approach resulted in >50% significant regressions, regardless of whether the test was applied directly after the expansion or much later when no signal of RE remained. This was also true, although to a lesser degree, in cases where the population was panmictic, particularly early in the simulations (Supplementary Fig. S2a). Since the power of linear regressions depend on the number of populations, there was no increase in the proportion of rejected null hypotheses with increasing number of cycles.

As indicated in Figure 2, when expansions started from d_1_, the lowest ψ and the highest heterozygosities were never found at the edge, even at the first time point after the expansions had finished (t_0_). This is reflected in the accuracy of both the TDoA and the non-TDoA, which is low from the beginning, and continues to decrease until the estimated origin coincides with the center of diversity caused by BEs (Supplementary Fig. S2b). When the expansion starts from the center, RMSE is low throughout the simulations (Supplementary Fig. S2b). Since the signal of a RE declines slower for ψ than genetic diversity, we also observed that the (negative) correlation between these two statistics started out strong, temporarily declined and again increased as the signal of the RE in ψ had diminished and both statistics were equally affected by BEs (Supplementary Fig. S3).

### Continuous space simulations in heterogeneous landscapes

In the continuous space simulations, the mean *F_ST_* between the two most distant populations at the end of the simulations were 0.12, 0.32, and 0. 62 (HL models) and 0.13, 0.40 and 0.73 (UL models) for high, medium and low levels of gene flow, respectively. The slightly higher levels of genetic differentiation in the UL models resulted from the populations in the HL models being connected by corridors of lower resistance along shallow areas along coasts (Fig. 1c).

The results from the continuous space simulations were concordant with the results observed in the stepping-stone models. For instance, following REs, the proportions of significant ψ first declined and then increased (Supplementary Fig. S4). The proportion of rejected null-hypotheses converged to similar values 2k-20k generations after the expansions started in both RE and DE models (depending on simulation parameter settings; Supplementary Fig. S4). The main difference is that the RE models started from a high proportion of rejected null-hypotheses (and then declined before increasing again), whereas the RE started from lower values and continually increased. When only 10k SNPs were sampled for these analyses and gene flow was low, there was not enough power in the binomial test to reject the null-hypothesis after the initial signal of RE had disappeared, even when geographic clines in ψ due BEs clearly existed, as was evident when using larger numbers of SNPs (50k; Supplementary Fig. S4).

As with the stepping stone models, *p*-values from TDoA analyses were highly inflated relative to non-TDoA results, where the latter could to some extent distinguish a range expansion from a demographic expansion or an equilibrium situation (with both ψ and Δ*_het_*) but only when gene flow was low (top panels in Supplementary Fig. S5 a,b). However, when gene flow was medium or high, the non-TDoA method could only reliably detect range expansions when the landscape was uniform (UL; left vs. right panels in Supplementary Fig. S5 a,b). In contrast to non-TDoA analyses, the proportion of significant correlations for the TDoA analyses was a positive function of time in both the RE and DE simulations suggesting BEs have an even stronger influence on TDoA than REs (Supplementary Fig. S5). A similar bias of estimated origins towards the meta- population center as seen for the stepping-stone models was observed for the continuous space models (based on the TDoA method; Supplementary Fig. S5 b; the estimated origins are shown in Supplementary Fig. S6). The correlation between ψ and *H_E_* was much stronger across time in the continuous space RE simulations (min *r^2^*∼0.6) compared to the stepping stone simulations (min *r^2^*∼0.25), especially when gene flow was high (min *r^2^*>0.8; Supplementary Fig. S3 and Supplementary Fig. S7). The correlation between ψ and *H_E_* was high across all levels of population connectivity in DE simulations (min *r^2^*>0.8).

### Predicting boundary effects at equilibrium in simulated data

Across all simulated data sets, *p*-values from the TdoA were inflated by a factor of 38 relative to *p*-values from the non-TdoA approach (in a quantile-quantile plot on a log_10_ scale). Despite this, the relationships between their effect sizes, *τ_TDoA_* and *τ_non_*_−*TDoA*_, respectively, were close to unity (Supplementary Fig. S8). However, in the empirical data, *τ_TDoA_* was inflated relative to *τ_non_*_−*TDoA*_ by a factor of 1.25, possibly due to more heterogeneous *N_e_*’s among natural populations affecting *τ_TDoA_* more than *τ_TDoA_*. We therefore used *τ*=*τ_non_*_−*TDoA*_ in the following analyses.

Below we considered 1D, 2D, and the continuous space simulations: RE-UL, RE-HL, DE-UL and DE-HL as different “simulation data sets”. Figure 3 illustrates the relationships between mean |ψ|, *F_ST_* and ɛ=|ψ|/*F_ST_* showing that for the simulation data sets where Res occurred (all but DE simulations), ɛ declines with time and reaches a background level where ɛ∼ɛ_eq_ (in DE simulations ɛ∼ɛ_eq_ throughout all simulation cycles). As also seen in Figure 3, ɛ normalizes the variance in |ψ| such that gene flow only explains <2% of the variation in ɛ_eq_ and instead most of the variation was found between the different simulation data sets (77%; Supplementary Fig. S9). Within each of them, some differences between levels of gene flow were also found as the interaction term between gene flow and data set explained 91% of the variation (Supplementary Fig. S9).

**Figure 3.**
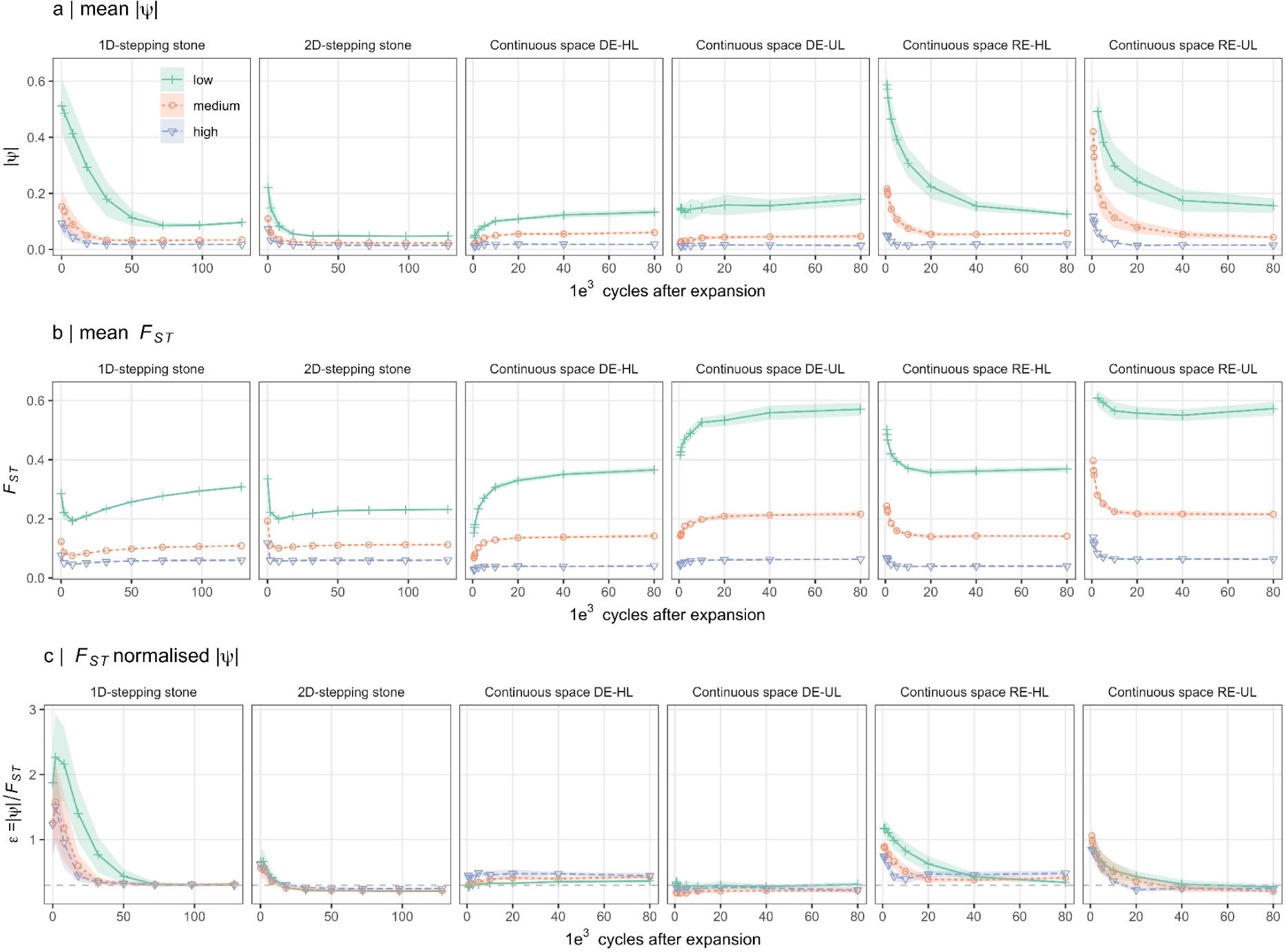
*F_ST_*normalized |ψ|. Shows the change of mean |ψ| (a), *F_ST_* (b) and ɛ=|ψ*|/F_ST_* (c) as a function of time in the simulations for three levels of gene flow; low (green), medium (red) and high (blue) for 1D and 2D stepping stone simulations as well as for the continuous space simulations with RE (RE) and when the expansion did not include a spatial component (demographic expansion, DE) in heterogeneous (HL) and uniform landscapes (UL). For 1D and 2D simulations the x-axis represents the number of cycles after the range expansion finished and for the continuous space simulations the x-axis shows the number of cycles after expansions started. In (c) the horizontal dashed line shows the mean ɛ at the end of the simulations representing the mean ɛ_eq_ (the value of ɛ at the end of each simulation replicate) across all the simulated data set (see also Supplementary Fig. S9).

In addition to the simulation data sets considered above, we further consider all data sets pooled (“All”), all data sets that include Res pooled (1D, 2D, RE-UL and RE-HL) and all data sets except the panmictic data (“Excl. panm.”) separately in the following analyses. Among these data sets, *τ_TDoA_* and |ψ| explained <40% (mean 15%) and <1%-89% (mean 48%) of the variation in *E*, respectively. However, when normalizing |ψ| with *F_ST_* (ɛ=|ψ|/*F_ST_*), the amount of variation explained increased to a minimum of 27% for all data sets (mean 65%) with 81%-95% of the variation in *E* explained in 4/7 data sets (Excl. panm., 2D, RE-UL and RE-HL). When considering each RE simulation data set independently (1D, 2D, RE-UL and RE-HL), 91%-99% of the variation in *E* could be explained by the full model (*E*∼ɛ\**τ*, equation 2) but when considering all data sets jointly (including the panmictic data), this number dropped to 78%. This is because of the slightly different background levels of ɛ_eq_ (see above) in the different simulations increasing the residual.

When excluding the panmictic data, ɛ explained 81% of the variation and the fit did not improve by adding τ to the model (Fig. 4) in contrast to the full data set where ɛ only explained 33%. This difference stems from low *F_ST_* in the panmictic data sets resulting in high ɛ but weak geographic clines in ψ. As range expansions require clinal variation in ψ, when predicting *E* from ψ, *F_ST_*and τ, the full model (*E*=1.00 + 0.0914ɛ - 0.894τ + 3.38ɛτ) fitted using all the simulated data was used, unless otherwise stated. We further considered data sets from RE models with ɛ/ɛ_eq_>1.2 as non-equilibrium data sets (true positives). Because of lower than average ɛ_eq_ (Fig. 3 and Supplementary Fig. S9), the power to detect Res in these data sets (the proportion of data sets exceeding the lower prediction boundary for *E*>1) was lower in 2D and RE-UL data sets (7% and 52%, respectively) compared to RE-HL data sets (86%). In our simulations, all data sets deriving from DE simulations as well as those sampled from the second half of the simulations (≤72k and ≤40k cycles for the stepping stone and continuous space models) are subsequently considered as equilibrium data sets (where ɛ∼ɛ_eq_). Among these, none exceeded the lower prediction boundary for *E*>1 (Figure 5).

**Figure 4.**
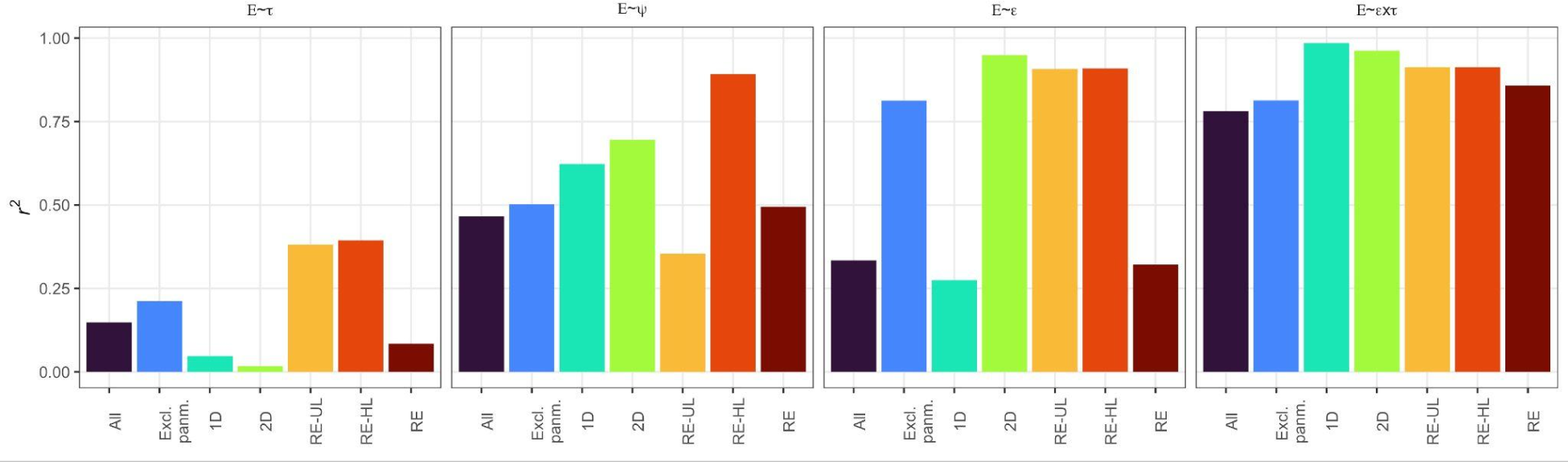
Predicting *E* from ψ, *F_ST_* and τ. Shows the proportion of variance explained (*r^2^*) when fitting the models *E∼*τ, *E∼*ψ, *E∼*ɛ and *E∼*ɛ*τ to different combinations of simulated data sets, where *E=*ɛ/*ɛ_eq_*, ɛ=|ψ|/*F_ST_* for a given data set and *ɛ_eq_* is the same value at equilibrium (for a given simulation replicate) and τ is the strongest positive *r^2^*for a linear regression between geographic distance and ψ for any population in the data (i.e. the most likely expansion origin). The data sets are: “All”-all data simulated data; “Excl. panm.”-all data except panmictic; “1D” - 1D stepping stone simulations; “2D” - 2D stepping stone simulations; “RE-UL” - continuous space simulations with uniform landscape and “RE-HL” - continuous space simulations with heterogeneous landscape and “RE” - all data sets where REs occurred (1D, 2D, RE-UL and RE- HL).

**Figure 5.**
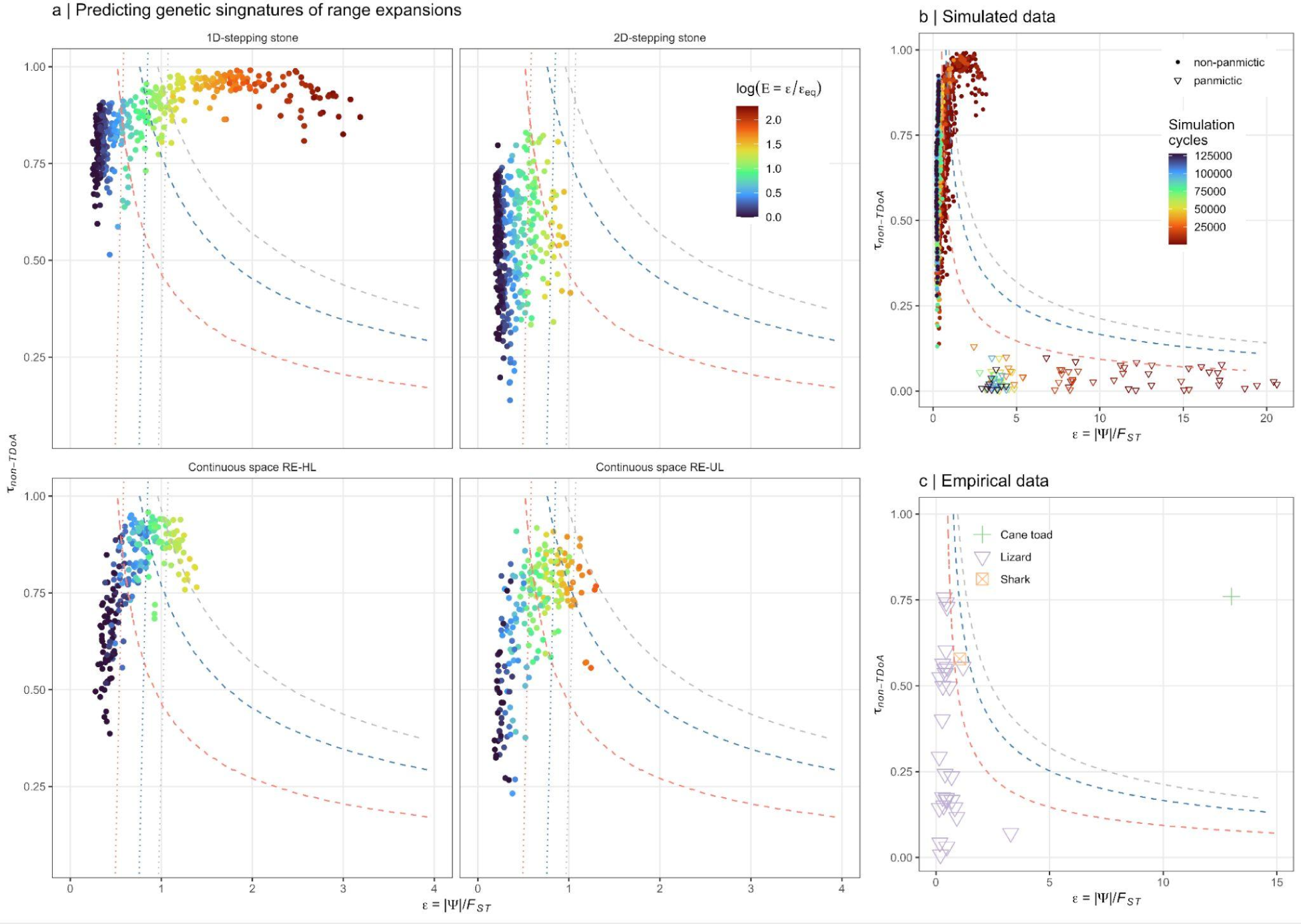
Predicting genetic signatures of range expansions. (a) Shows the effect size (τ) from non-TDoA analyses (the strongest positive *r^2^* for a linear regression between geographic distance and ψ for any population in the data) as a function of ɛ=|ψ|/*F_ST_.* Color represents *E=*ɛ/*ɛ_eq_*in the simulated data (where ɛ_eq_ can be known) on a log scale and the dashed lines indicate the lower limits of the prediction interval for *E*>1 for α=0.05 (red) α=0.05/30 (blue; the Bonferroni corrected significance level for the empirical data) and α=0.0001 (grey) when fitting *E∼e\**τ to data set “All” (see Fig. 4) explaining 77% of the variation in the data. The dotted lines are the same lower boundary limits as above except for a model fitted for data set “Excl. Panm.” (see Fig. 4) that explains 81% of the variation in the data. (b) shows the same data as in (a) but also including the panmictic data sets (triangles; *F_ST_* fixed at 0.001) where color represents the number of cycles since range expansion finished (stepping-stone models) or since range expansion started (continuous space simulations). In (c) results from 30 empirical data sets from scincid lizards (violet, n=28), blacktip shark (orange, n=1) and the invasive cane toad (green, n=1) are shown.

The mean effect sizes for TdoA and non-TdoA were *τ_TDoA_*=0.47 and *τ_non_*_−_*_TDoA_*=0.45 for equilibrium data sets and *τ_TDoA_*=0.71and *τ_non_*_−_*_TDoA_*=0.76 for non-equilibrium data sets, respectively and the mean *ɛ* was 2.2x higher in the non-equilibrium data sets (*ɛ* =0.69) compared to equilibrium data sets (*ɛ* =0.32).

### Is there any evidence for range expansions in the empirical data?

Population structuring was generally high among the 28 Australian scincid lizards datasets ranging from *F_ST_*=0.10 to *F_ST_*=0.72 between the two most differentiated populations, covering a similar range of values as our simulated data. Among these, a minimum of 33% significant pairwise ψ-values were found for all data sets, ranging up to 100% (mean=65%). Significant clines in ψ were found in 27/28 data sets with TdoA method and in 14/28 data sets with non- TdoA method. Notably, however, many of these data sets had weaker clinal variation in ψ (*τ_TDoA_*=0.55 and *τ_non_*_−*TDoA*_=0.35) than in the equilibrium data sets from the simulations (see above). In addition, the data sets with high ɛ tended to be those with the lowest *τ* (Fig. 5c) and consequently, while only a single data point exceeded the lower boundary for *E*>1 at the 95% confidence limit, no tests were significant after accounting for multiple testing (dashed lines in Fig. 5). Furthermore, these empirical data sets tended to cluster more with equilibrium data sets, except for one data set where ɛ=3.3. However, for this data the geographic cline in ψ was also weak (*τ_non_*_−*TDoA*_=0.071) and this was also the data set with the lowest level of genetic differentiation. The high ɛ in this data point could thus have resulted from a combination of low *F_ST_* and relatively high (given the level of population structuring), but non-clinal, variation in *N_e_*. Notably, if panmictic data were not included in the model fitting, four of the data sets exceed the Bonferroni corrected lower boundary for *E*>1 (dotted lines in Fig. 5) due to their relatively high ɛ. In the blacktip shark data *F_ST_*=0.77 between the two most differentiated populations. The scaled ψ for this data set was estimated to ɛ=1.05 and the strongest correlation between geographic distance and ψ was estimated to *τ_non_*_−*TDoA*_=0.58 resulting in an estimated *E*=2.63 which only exceeded the lower prediction boundary for *E*>1 when not correcting for multiple testing (Fig. 5 c).

In contrast to the above data sets, the genetic structuring among the cane toad populations was much lower (*F_ST_*=0.039 between the two most differentiated populations). Since the mean *F_ST_*<0 between all pairwise comparisons, as for the panmictic data from the simulations *F_ST_*=0.001 was used to calculate ɛ. With |ψ|=0.0134, ɛ=13.4 and *τ_non_*_−*TDoA*_=¿0.76, *E* was estimated to be 4.5. Such high predicted values for *E* were only observed in the 1D-stepping stone simulations where the difference between equilibrium and non-equilibrium data sets were the highest (*E_max_*=9.9; *E_max_*=4.2 for any of the other simulations). Consistent with the historical records of a rapid RE from Gordonvale, North Queensland since its introduction in 1935, the estimated origin using TdoA was highly accurate for both ψ and Δ*_het_*(Supplementary Fig. S10).

## Discussion

Using individual based spatially explicit forward-in-time simulations we demonstrate that ψ and *H_E_* are similarly affected by Bes under mutation-drift and gene flow equilibrium scenarios, resulting in clines of decreasing diversity and increasing ψ from the meta-population center towards the edges. This is because the same processes that lead to Bes in genetic diversity (Wilkins and Wakeley 2002) also cause asymmetries in the SFS (Gutenkunst et al. 2009). As a consequence, not knowing a priori the underlying population demographic model and level of connectivity, it is not possible to specify a threshold value of ψ that can reliably differentiate a RE from a BE in population genomic data. This is particularly true since we show that detecting significant asymmetries in 2D-SFS is only a matter of statistical power and thus, in contrast to what was previously proposed by Peter and Slaktin (2013, 2015), rejecting the null-hypothesis of ψ≠0 cannot be taken as evidence of Res even when there also are significant geographic clines in ψ. Thus, the relevant question is not whether significant asymmetries in the SFS exist, but rather whether they are stronger than can be expected due to Bes under a specific population demographic scenario and level of population connectivity.

Our results suggest we should be careful when interpreting the results from recent population genetic studies that used clines of genetic diversity and ψ to test for Res and identify their centers of origin. For example, Maisano-Delser et al. (2019), Walsh et al. (2022) and Lesturgie et al. (2023) identified range expansions of coral-reef associated sharks originating from the Malay Archipelago and speculated on the potential role of this region as a refugium for coral-reef associated organisms from which recolonization started. Given that we could not reject the null-hypothesis of *E*≤1 (after corrections for multiple testing) in the blacktip shark data, and the fact that the Malay archipelago is located close to the center of the distribution of Indo-Pacific coral-reef associate sharks, indicates that the observed geographic patterns in genetic diversity and ψ could also have been caused by Bes. Similarly, we show that previous work that identified the center of origin of several lizard species in Australia was biased both by the limitation of these methods as well as by a coding error in the R package that implemented them (which resulted in ψ-matrices with inverted polarities), leading to the incorrect conclusion that many Australian lizard species had a center of origin at the periphery of their range. The same bug has likely affected the results of several other papers as well, such as Jaya et al. (2022), where an origin of range expansion was estimated to be at the very edge of the distribution range despite the fact that the highest levels of genetic diversity was seen at the center, and He et al. (2017), where TDoA performed suspiciously poorly in simulated data (see also below).

Consistent with the literature (Ioannidis et al. 2021; He, Prado, and Knowles 2017; Peter and Slatkin 2015, 2013), ψ and clines thereof were more sensitive to REs than statistics based on genetic diversity, yet in most cases they remained highly correlated in equilibrium and non- equilibrium meta-populations alike. Nevertheless, how these statistics have been utilized and interpreted in the context of REs in various studies has varied dramatically, with no clear consensus. Because no readily available software or R-function exists for rejecting the null- hypothesis of ψ≠0, most previous studies have not attempted to do so (but see data availability for access to the updated R-functions used in this study). Instead, the geographic patterns in the magnitude of ψ (or similar statistics that reflect asymmetries in SFS) have been interpreted in relation to other summary statistics (e.g. *H_E_*, Tajima’s D and IBD and overall patterns of genetic structuring) and independent sources of information regarding range expansions, such as historical records (Jaya et al. 2022; Ioannidis et al. 2021; Mestre et al. 2022; Pierce et al. 2014; Bringloe et al. 2022; Hemstrom et al. 2022).

While many of the limitations of ψ are often discussed (Ioannidis et al. 2021; He, Prado, and Knowles 2017; Mestre et al. 2022; Riginos et al. 2016), the possibility of BEs causing high false positive rates when testing for REs has, until now, been largely ignored. More importantly, while the effects of BEs on spatial patterns of ψ were reported in Peter and Slatkin (2013; 2015), the problem of BEs, as demonstrated here, appears to be much more severe than originally claimed. Even in a much more recent simulation study used to compare the performance of TDoA with the ABC based X-ORIGIN, the problems of BEs were not detected since the method was never evaluated under equilibrium conditions (He et al. 2017).

While the TDoA approach was never intended as a stand-alone test for REs in population genomic data, it has nevertheless been used to support REs, sometimes even without first formally rejecting the null-hypothesis of ψ≠0 (Maisano Delser et al. 2019; Singhal, Wrath, and Rabosky 2022; Peter and Slatkin 2015; Jaya et al. 2022; He, Prado, and Knowles 2017). Here we show that the *p*-values from TDoA are in practice meaningless since this test was significant at α=1e-07 in ∼50% of the panmictic data, where no clines in ψ could exist. While the TDoA approach can infer an origin also in unsampled geographic regions, we have here shown that this estimate can only be accurate immediately after a range expansion has finished, or when the origin already coincides with the center of the species range. We also urge caution when the estimated origin is inferred to be outside the convex hull of the sampling locations (at least using the implementation of TDoA available from the rangeExpansion R-package), since in the panmictic data sets the origin was estimated to be at d_1_ or d_81_ (the two most extreme demes in a 1D stepping stone chain) in 22% of the data sets instead of the expected 2% (assuming the estimated origin is randomly distributed among all 81 demes). The most reliable of the explicit tests for REs explored here was thus the more conservative non-TDoA approach that showed reasonable power to detect true REs with no false positive, but only in the RE-UL simulations or in the RE-HL simulations when gene flow was low; in all other simulations (except the panmictic data) ≫5% significant tests were observed in equilibrium populations as well as in simulated data where no REs occurred. The method X-ORIGIN (He, Prado, and Knowles 2017) has thus potentially several advantages over the TDoA and non-TDoA methods tested here - since it is based on ABC framework and relies on simulations, any influence of BEs are expected to be reflected in the uncertainties associated with the point estimates from this method, equilibrium data sets are also included.

The high generality of the model *E*∼ɛ*τ across a wide range of simulation approaches and parameter settings as well as its robustness against false positives in the simulated data opens up the possibility to also predict and test for *E*>1 in empirical data sets. This relies on the premise that the true ɛ_eq_ in empirical data has a similar distribution as in the simulated data here. The upper 95% quantiles of the distribution of ɛ_eq_ among all the simulated data was ɛ_eq_=0.48. Even though a minimum of 33% significant ψ values were observed for each of the empirical lizard data sets, all except one also with significant clines in ψ (based on the TDoA approach), ɛ exceeded 0.48 in only seven (out of 28) of these data sets. Notably, the upper 95% quantile for τ_non-TDoA_ for all equilibrium data sets from the simulations was 0.75 while the maximum observed τ_non-TDoA_ among the lizard data sets was 0.55. Thus, the levels of |ψ|, *F_ST_* and τ_non-TDoA_ observed in the empirical data are indistinguishable from those observed among the equilibrium data sets and consequently, none of the predicted *E*’s for these data sets exceeded the lower prediction boundary for *E*>1 (after accounting for multiple testing). Since the null hypothesis of *E*≤1 could not be rejected in any of the empirical lizard data sets the possibility of false positives is not a concern. It is, however, possible that the BEs in our simulations were exacerbated relative to the lizard populations, resulting in low statistical power to reject *E*≤1. Since both asymmetric gene flow as well as non-clinal variation in *N_e_* are expected to increase ɛ_eq_, both of which are likely to be common in natural populations but were not considered here, we regard this as highly unlikely. We are also not accounting for the fact that habitat quality may decrease towards the edges of a species distribution in natural populations (Sexton et al. 2009; Gaston 2009), increasing the clinal variation in *N_e_* beyond what can be expected by BEs alone. More generally, whenever habitat quality affects local population densities (and thus levels of genetic drift) and varies predictably in space, the highest diversity and lowest ψ are expected where the habitat quality is the highest, something that also needs to be considered when interpreting spatial patterns in ψ. In contrast to the lizard data sets, the null hypothesis that *E*≤1 could clearly be rejected in the cane toad data set, with an estimated ɛ=13.4 that is >4x higher than for any of the simulated data sets (ɛ_max_=3.18) and strong clinal variation in both ψ and *H_E_*. This is entirely consistent with the recent introduction (1935) in Gordonvale, North Queensland followed by a westward range expansion.

Using simulated data to predict signatures of REs in empirical data is predominantly presented here as an example of what can still be done, when the true null distribution (ɛ_eq_) in natural populations cannot be known. For instance, the number of replicates in our simulated data for each parameter combination was only n=10 but we still used all data sampled at multiple different time points from the simulations as data points in the models. Furthermore, we did not include any simulation with non-clinal variation in *N_e_,* asymmetric migration rates nor declining habitat quality toward the range boundaries. However, understanding the range of ɛ and τ that can be generated under equilibrium scenarios in the simulations can nevertheless greatly help us to evaluate how likely a true signal of REs exists in an empirical data set. For example, in the analyses of the five populations of *Arabidopsis thaliana* in Peter and Slatkin (2015) used to demonstrate the merits of ψ-based test for REs, the *p*-values reported for τ_TDoA_ ranged between 6.8e-6 and 4.2e-55, but the effect sizes for this test (τ_TDoA_) ranged only between 0.12 and 0.26. Such weak clines in ψ were observed in 78% of our equilibrium data sets and in <5% of our non- equilibrium data sets.

In conclusion, there are several practical implications emerging from our study. First, it is clear that although a strong correlation between geographic distance and ψ is expected from REs, the strengths of ψ-clines alone is not sufficient to determine whether a RE occurred. Second, the upper 95% quantile for ɛ_eq_ in our simulated data (ɛ_eq_=0.48) suggest that strong clinal variation in ψ is likely to reflect a true signature of REs only if the overall strength of ψ also is at least 50% of the mean pairwise *F_ST_* in the data. Third, predicting *E* only requires knowing |ψ|, *F_ST_* and τ_non-TDoA_, which can easily be estimated from population genomic data from natural populations and using this approach, our analyses suggest that the initial signatures of REs detected in the 28 lizard data sets and in the blacktip shark data set (using only ψ and clines thereof), likely are false positives. In the cane toad data, however, our analyses were highly concordant with the known history of a range expansion in this species, indicating that geographic clines in both ψ and *H_E_*can indeed be informative of REs provided the effects of BEs are accounted for.

## Data and Resource Availability

All data sets and analysis pipelines will be made available on data dryad once the manuscript has been accepted for publication.

## Acknowledgements

This work was funded by the University of Hong Kong. We would like to thank Sonal Singhal for providing the scincid lizards data and Sonal Singhal, Vince Buffalo and Caroline Dahms for valuable comments on the manuscript.

