## Supplementary data for "Boundary effects cause false signals of range expansions in population genomic data"

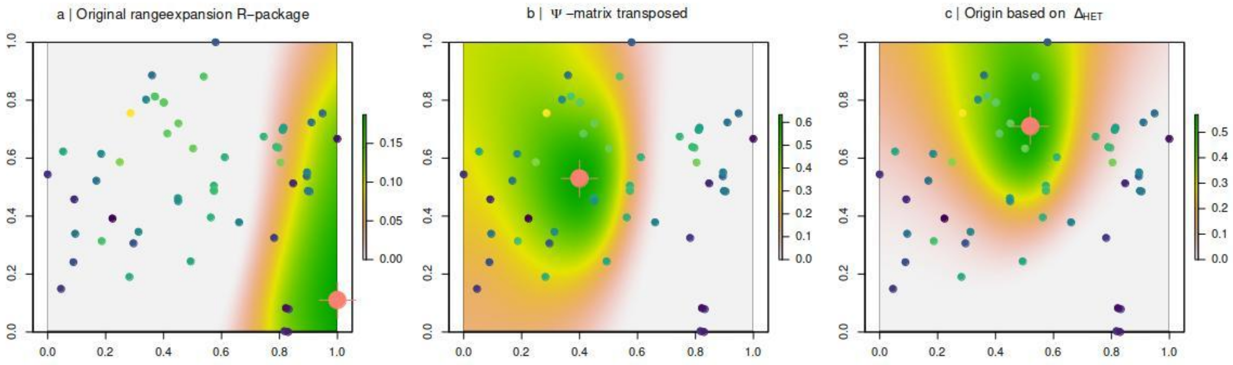


**Supplementary File S1 | Effect of a major bug fix in R-package rangeExpansion**. When first using the rangeExpansion R-package (v.0.0.0.9000) on our simulated data our results appeared to be opposite to what we expected; the estimated origin was always indicated for the population that was the last to be colonised. We traced this back to a bug in lines 320 and 321 for the function *get.all.psi* in the file; https://github.com/BenjaminPeter/rangeexpansion/blob/master/R/re_functions2.r:

```
mat[j,i] <- get.psi( ni, nj, fi, fj,resampling=resampling, n=n )
mat[i,j] <- -mat[j,i]
```

where the indexes for rows (i) and columns (j) had been reversed, ultimately resulting in a transposed ψ-matrix. In the above figure we show the results of the TDoA approach for an empirical lizard data set (*Ctenotus inornatus*) when using the original code (a) and the same analyses with the polarity of the original ψ-matrix inverted (using our modified function *get.all.psi.mc.bin*). The color of the dots are coordinates for the populations with color indicating genetic diversity ($H_E$) and the color of the background represents the effect sizes from TDoA analyses ($r^2$) with the estimated origin indicated by red.

Figure (c) shows the TDoA analyses using the difference in heterozygosities between two populations ($\Delta_{het}$) instead of ψ, where the origin is expected to have a negative correlation between geographic distance and $\Delta_{het}$ (genetic variation decreases away from the estimated origin). This agrees much better with the analyses based on the correct ψ-matrix but more importantly using the correct orientation of the ψ-matrix gives expected results across all the simulated data (see for instance Fig. 2, main document). See also Supplementary Figure S10 for an example using the cane toad data.

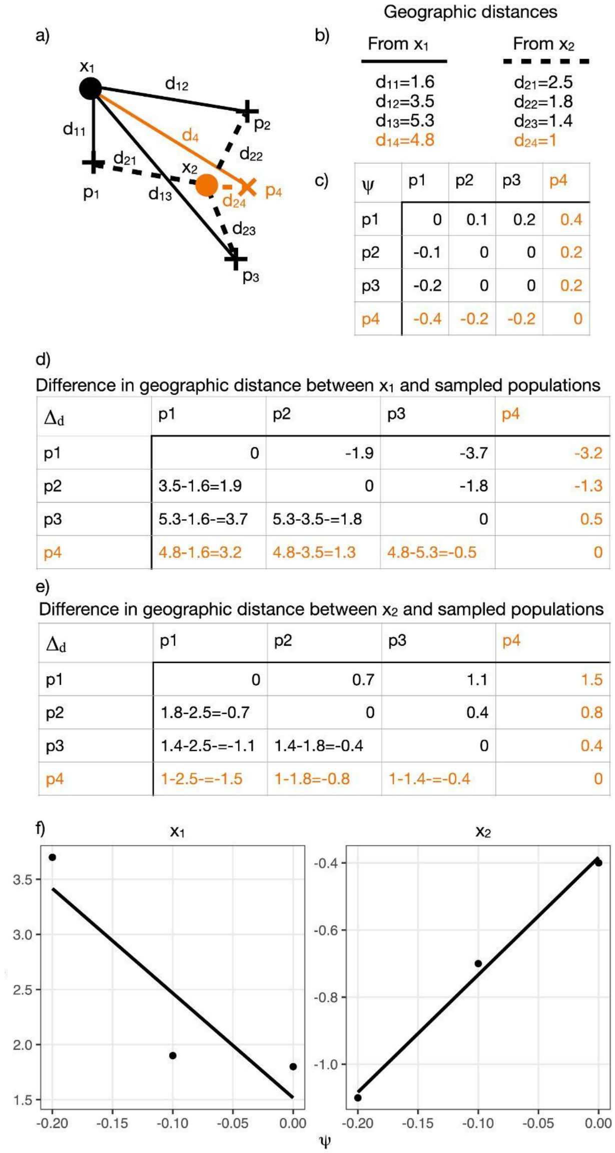


c)

| ψ | p1 | p2 | p3 | p4 |
|---|---|---|---|---|
| p1 | 0 | 0.1 | 0.2 | 0.4 |
| p2 | -0.1 | 0 | 0 | 0.2 |
| p3 | -0.2 | 0 | 0 | 0.2 |
| p4 | -0.4 | -0.2 | -0.2 | 0 |

d) Difference in geographic distance between $x_1$ and sampled populations

| $\Delta_d$ | p1 | p2 | p3 | p4 |
|---|---|---|---|---|
| p1 | 0 | -1.9 | -3.7 | -3.2 |
| p2 | 3.5-1.6=1.9 | 0 | -1.8 | -1.3 |
| p3 | 5.3-1.6-=3.7 | 5.3-3.5-=1.8 | 0 | 0.5 |
| p4 | 4.8-1.6=3.2 | 4.8-3.5=1.3 | 4.8-5.3=-0.5 | 0 |

e) Difference in geographic distance between $x_2$ and sampled populations

| $\Delta_d$ | p1 | p2 | p3 | p4 |
|---|---|---|---|---|
| p1 | 0 | 0.7 | 1.1 | 1.5 |
| p2 | 1.8-2.5=-0.7 | 0 | 0.4 | 0.8 |
| p3 | 1.4-2.5-=-1.1 | 1.4-1.8=-0.4 | 0 | 0.4 |
| p4 | 1-2.5-=-1.5 | 1-1.8=-0.8 | 1-1.4-=-0.4 | 0 |

**Supplementary File S2 | Implementation of Time Difference of Arrival (TDoA) in the rangeExpansin R-package.** In the figure (left) we demonstrate the TDoA approach as described in Peter and Slatkin (2013; 2015) and implemented in the rangeExpansion R-package. While the original implementation of the TDoA approach relies on finding a set of points that has the same difference in distance to the origin corresponding to the arm of a hyperbola (see Fig. 4 in Peter and Slatking, 2013), the function *prep.tdoa.data* evaluated this equation on a grid and finds the location with the highest correlation corresponding the origin of the expansion. In (a) $x_1$ and $x_2$ represent two alternative grid points and $p_1$-$p_4$ represent population samples (shown as "+"), of which $p_4$ indicated in orange ("x") is the true origin of the range expansion. The distances from $x_1$ to $p_1$-$p_4$ (solid line) and from $x_2$ to $p_1$-$p_4$ (dashed line) are shown in (b). The pairwise ψ-matrix is shown in (c) and the difference in distance ($\Delta_d$) between a grid point and each pair of populations for a given entry in the ψ-matrix for $x_1$ and $x_2$ are shown in (d) and (e). The correlation between the pairwise ψ-values and the corresponding $\Delta_d$ values for $x_1$ and $x_2$ are shown in (f) showing that since $x_2$ (indicated in orange in a) has the strongest positive correlation it is located closest to the true origin. Note that since the correlation in (f) is based on two pairwise distance matrices (c vs. d or c vs f) the significance of the relationship between geographic distance and ψ should be evaluated e.g. by a mantel test.

Peter BM, Slatkin M. 2013. Detecting range expansions from genetic data. Evolution 67:3274–3289.
Peter BM, Slatkin M. 2015. The effective founder effect in a spatially expanding population. Evolution 69:721–734.

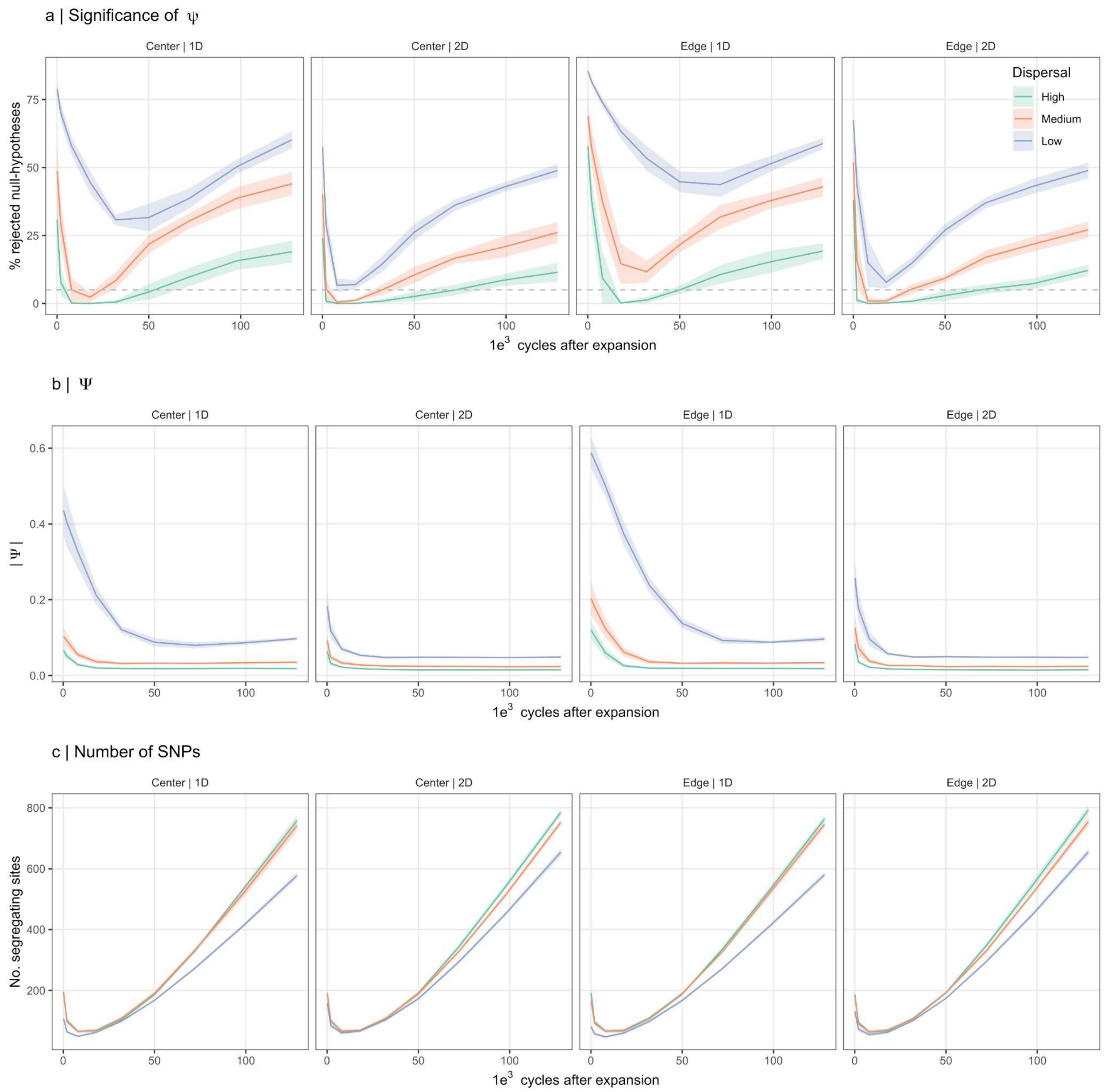


**Supplementary Figure S1 | Significance of ψ increases with time due to increased statistical power.** This figure shows the proportion of rejected null-hypotheses as a function of time since range expansions ended for ψ≠0 for data from the stepping stone simulations (a), the mean absolute ψ (b) and the mean number of segregating sites in each 2D-SFS used to test significance (c). Simulations with high, medium and low levels of gene flow are shown in green, red and blue, respectively. The pattern of declining proportions of rejected null-hypotheses followed by an increase in (a) is not due to corresponding changes in ψ (b) but rather an increase in the number of SNPs used for testing the significance of ψ (c) i.e., statistical power. Dashed horizontal line (a) indicates the expected number of significant tests under the null-hypothesis for $\alpha$=0.05. Shaded area indicates standard deviation.

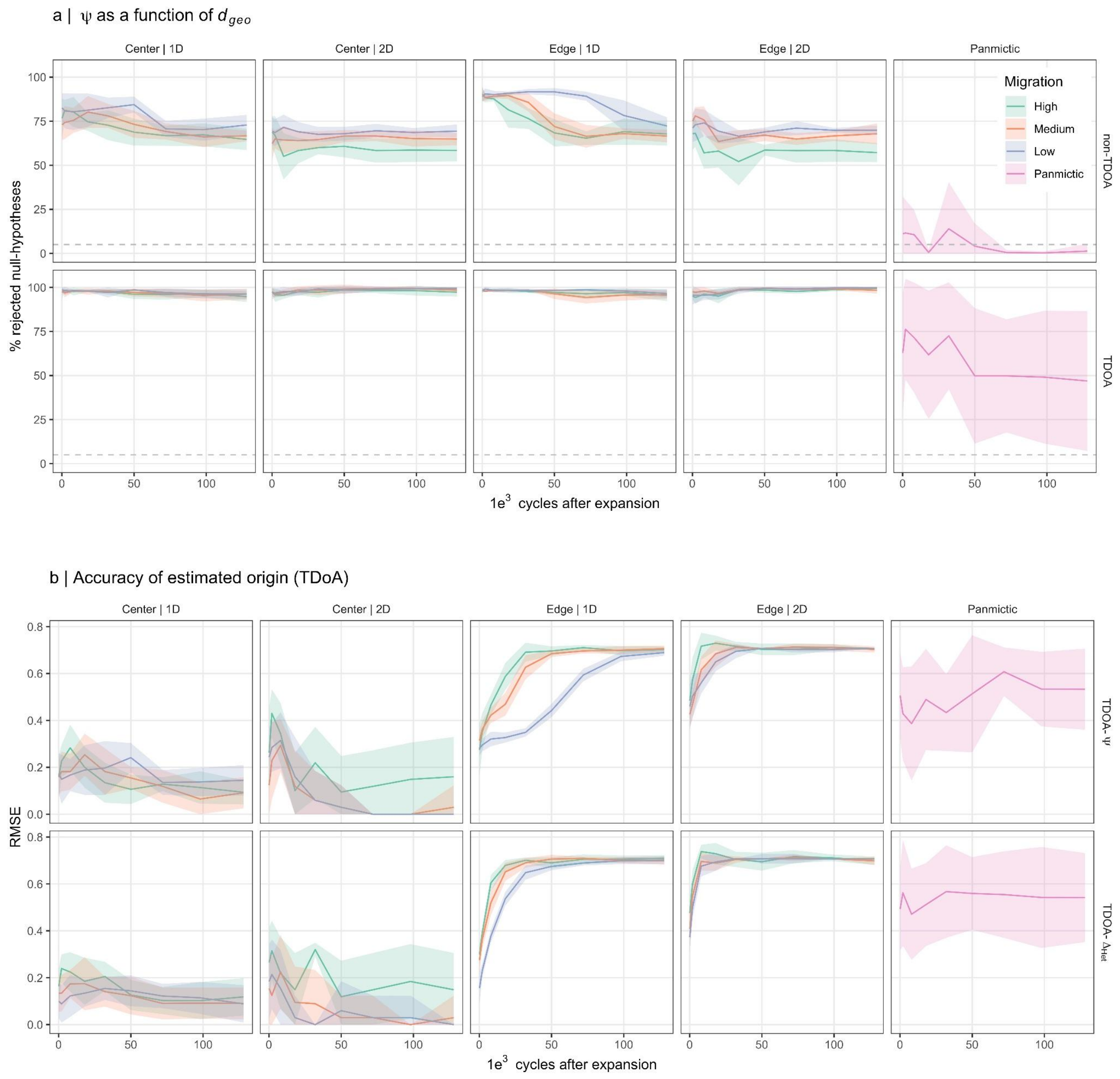


**Supplementary Figure S2 | Stepping stone simulations - ψ and $H_E$ as a function of geographic distance**. The proportion of significant correlations between ψ or $H_E$ and geographic distance are shown in (a) for the non-TDoA (top panel) and TDoA approaches (bottom panel) as a function of time since range expansions ended. When using the TDoA approach to estimate the origin of range expansions (b), The accuracy (RMSE) remains low throughout the simulations when the expansion originated from the center, but increases rapidly for expansions starting from the edge. This decline, however, is faster for $H_E$ than for ψ, particularly in the 1D simulations (top panel versus bottom panel for facet “Edge | 1D”). Dashed horizontal line (a) indicates the expected number of significant tests under the null-hypothesis for $\alpha$=0.05. Shaded area indicates standard deviation.

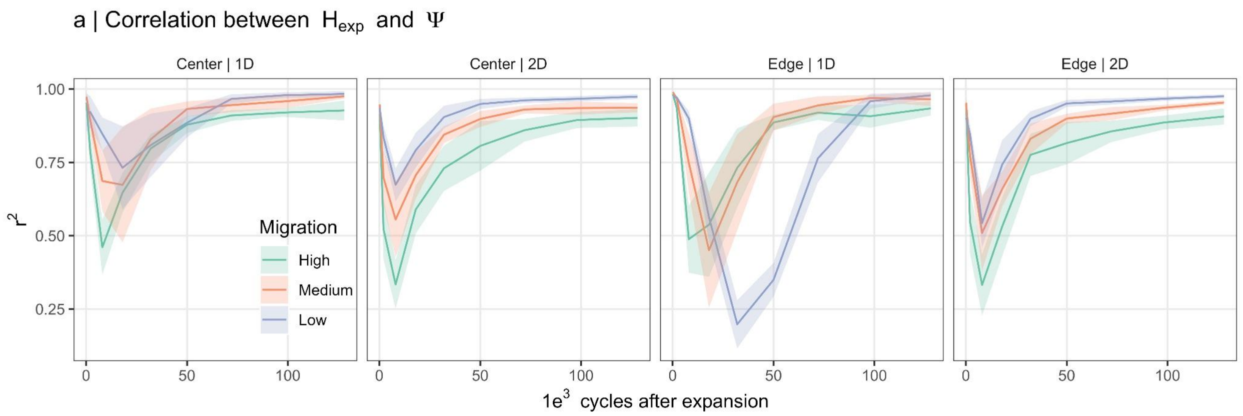


**Supplementary Figure S3 | Correlation between genetic diversity and ψ in stepping stone simulations**. Directly after the range expansions ended, the correlation between $\Delta_{het}$ and ψ was generally strong ($r^2$~1), but since the signature of range expansions decline faster for $H_E$ than for ψ, this correlation initially decreases with time and then increases again as the effects of the initial range expansions declines and the data instead become more affected by boundary effects. Shaded area indicates standard deviation.

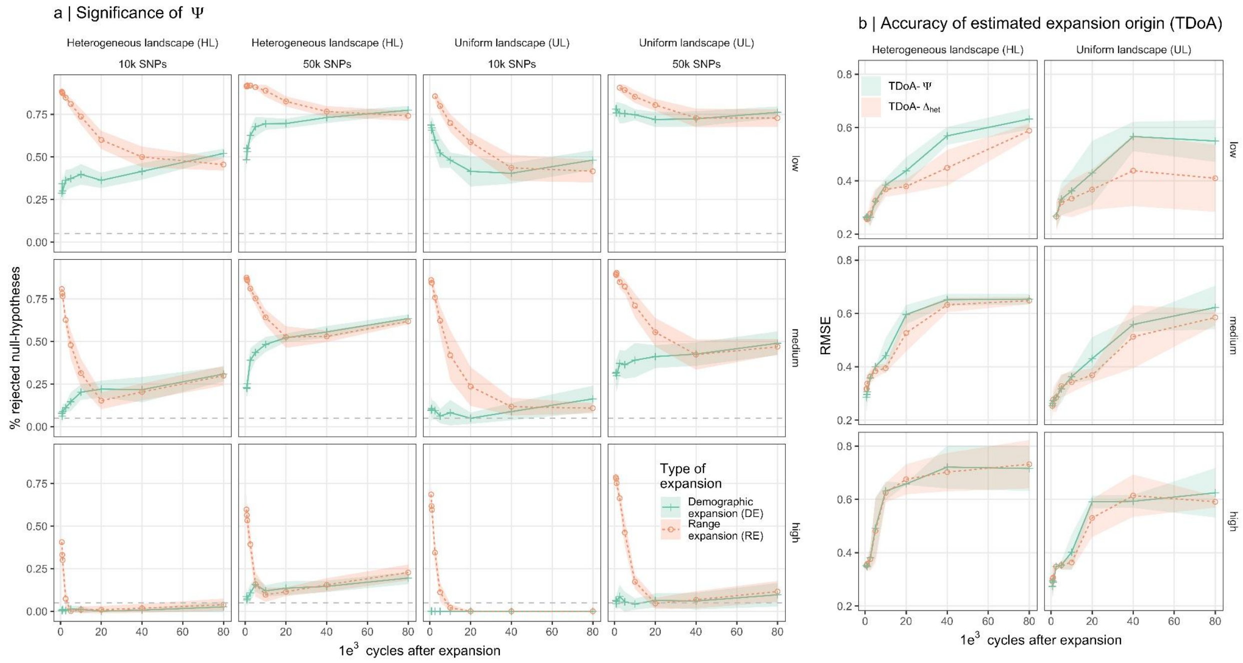


**Supplementary Figure S4 | Continuous space simulations**. Shows the proportion of rejected null-hypotheses for ψ≠0 for the continuous space simulations (a) and the accuracy of the TDoA approach used to estimate the origins of range expansions (b) for low, medium and high dispersal (top to bottom). Since the power of the binomial test (used in a), depends on the number of SNPs segregating in both populations in 2D-SFS, no false positive tests were detected for simulations when gene flow was high using a 10k random subset of the SNPs in the analyses. In all other cases, ≫5% of significant tests (above dashed horizontal line) were found also for simulations where the demographic expansion lacked a spatial component (DE). In contrast to the RE simulations, the proportions of significant tests increases with time in the DE simulations such that both approach similar values after the initial signal of range expansions in RE has disappeared. The RMSE is scaled such that RMSE=1 for two points at the opposite corners of the map. Since the true origin was at the top left corner of the map (Fig. 1e, main text), the accuracy of TDoA estimated range expansion origins rapidly declined after the range expansions started as the patterns in the data become increasingly dominated by boundary effects (b; see also Supplementary Fig. S6).

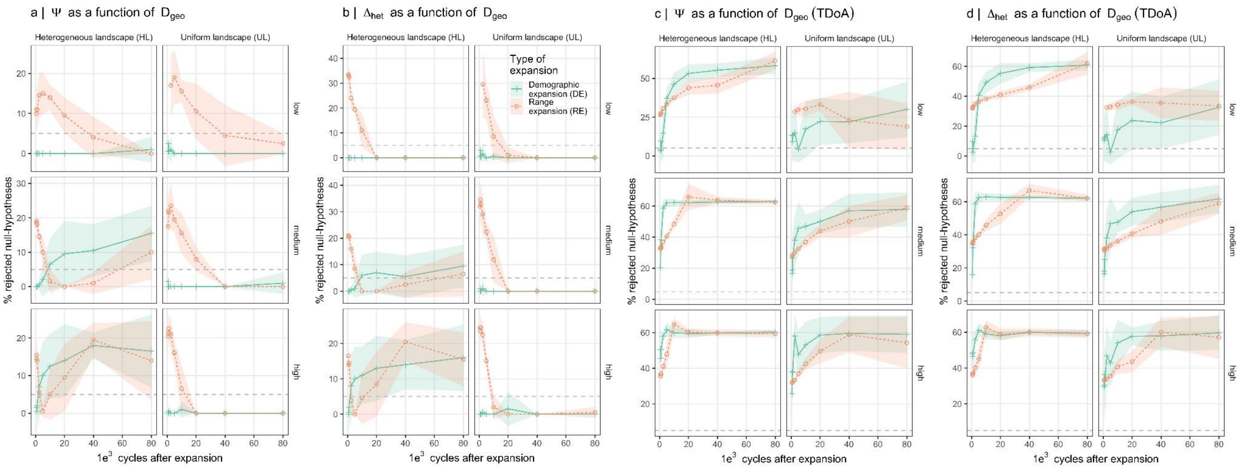


**Supplementary Figure S5 | Continuous space simulations - ψ and $H_E$ as a function of geographic distance.** Shows the proportion of significant correlations in non-TDoA (a,b) and TDoA analyses (c,d) for the continuous space simulations. Using the more conservative non-TDoA shows that, for both ψ and $H_E$, when the landscape is uniform (UL), <5% significant tests are found when a demographic expansions lacks a spatial component (DE, green), and >5% significant test are only found when a true signal of range expansion existed in the data (red, compare for instance with Supplementary Fig. S1 b). However, when the landscape is heterogeneous (HL) $\gg$5% significant tests are produced also for UL simulations, except when gene flow is low. In contrast, the proportion of significant correlations for the TDoA approaches is predominantly a positive function of time (simulation cycles) for both RE and DE simulations indicating that boundary effects have an even stronger effect on TDoA than range expansions.

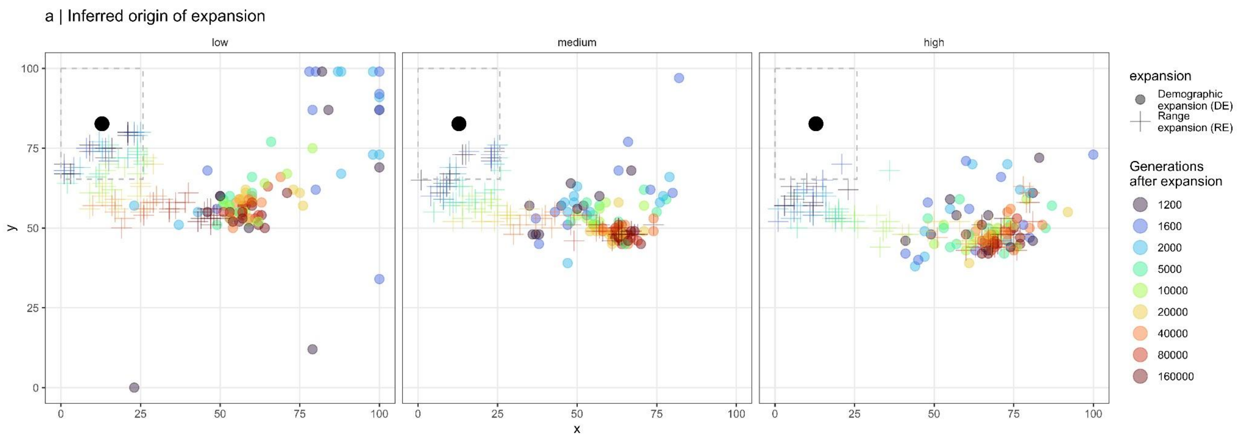


**Supplementary Figure S6 | Estimated origins based on TDoA for continuous space simulations**. Shows the estimated origins using the TDoA approach for continuous space simulations when the landscape is heterogeneous. Color indicates no. cycles after expansion started with dots indicating demographic expansions without a spatial component (DE) and range expansion and “+” indicates simulations with range expansion (RE). The box indicated by dashed line in the top left corner indicates the equilibrium population range before expansion started and the black dot indicates the center of this box that marks the position of the true origin used for analyses in Supplementary Figure S4.

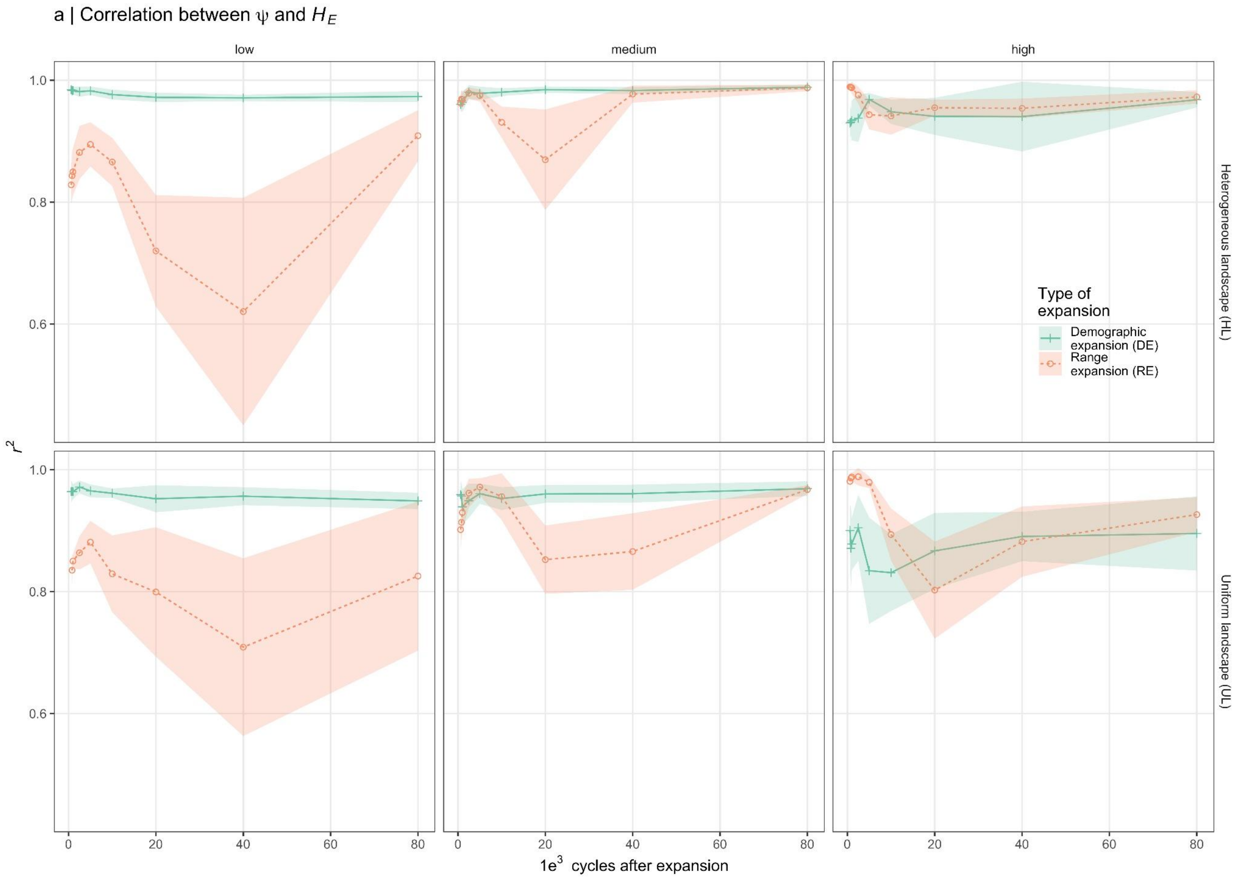


**Supplementary Figure S7 | Correlation between genetic diversity and ψ in continuous space simulations**. In contrast to the stepping stone simulations, where, during a transitional period of declining correlations between $\Delta_{het}$ and ψ (Supplementary Fig. S3) in the continuous space simulations $\Delta_{het}$ and ψ remain highly correlated ($r^2$≨0.8) with time, regardless of whether demographic expansions contain a spatial component or not (DE vs. RE simulations, respectively), except when dispersal is low (left) or when dispersal is high and the landscape is uniform (UL; bottom right).

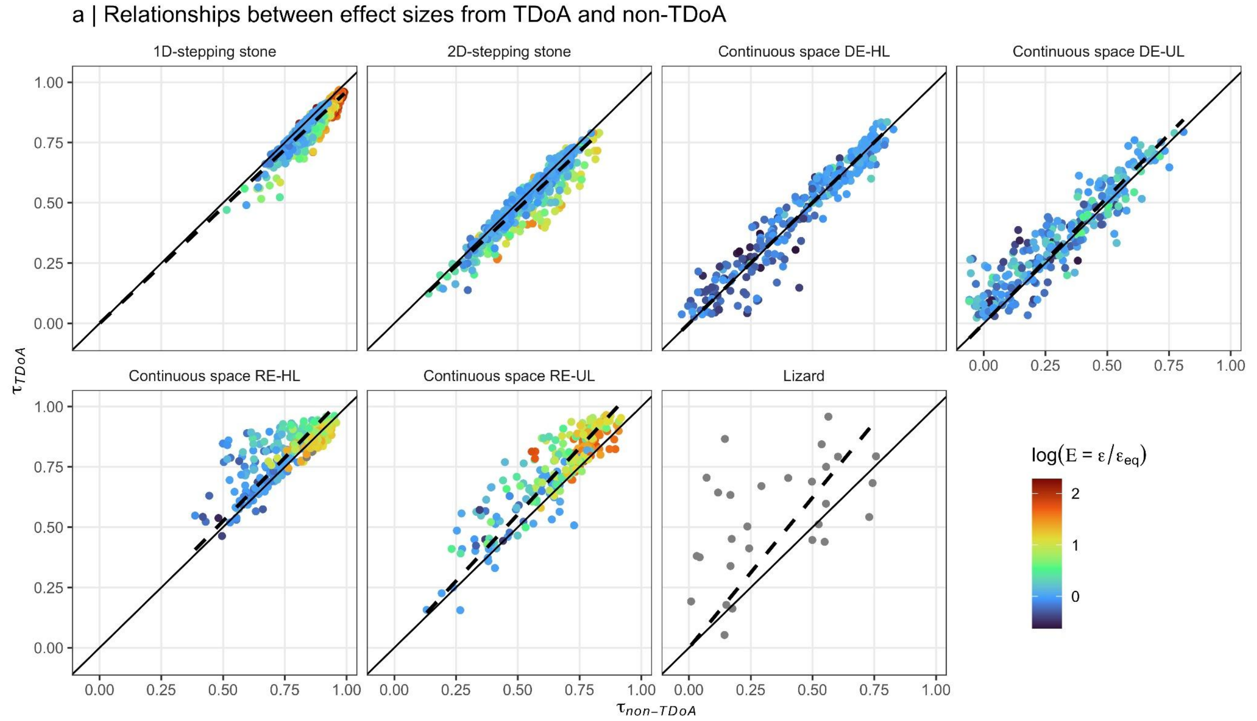


**Supplementary Figure S8 | The relationships between effect sizes ($\tau$) from TDoA and non-TDoA analyses**. The effect size ($\tau$) from TDoA and non-TDoA is the $r^2$ for the population or grid point, respectively, with the strongest positive correlation between geographic distance and ψ. Despite *p*-values from the TDoA being highly inflated relative to *p*-values from the non-TDoA approach, the relationship between their effect sizes were close to unity in the simulation data sets ($r^2$ ranging between 0.98 and 1.00 and slope ranging between 0.95 and 1.12). However among the empirical lizard data sets values from TDoA tended to be overestimated relative to non-TDoA (slope=1.25 $r^2$=0.77). Color shows E estimated from the simulated data.

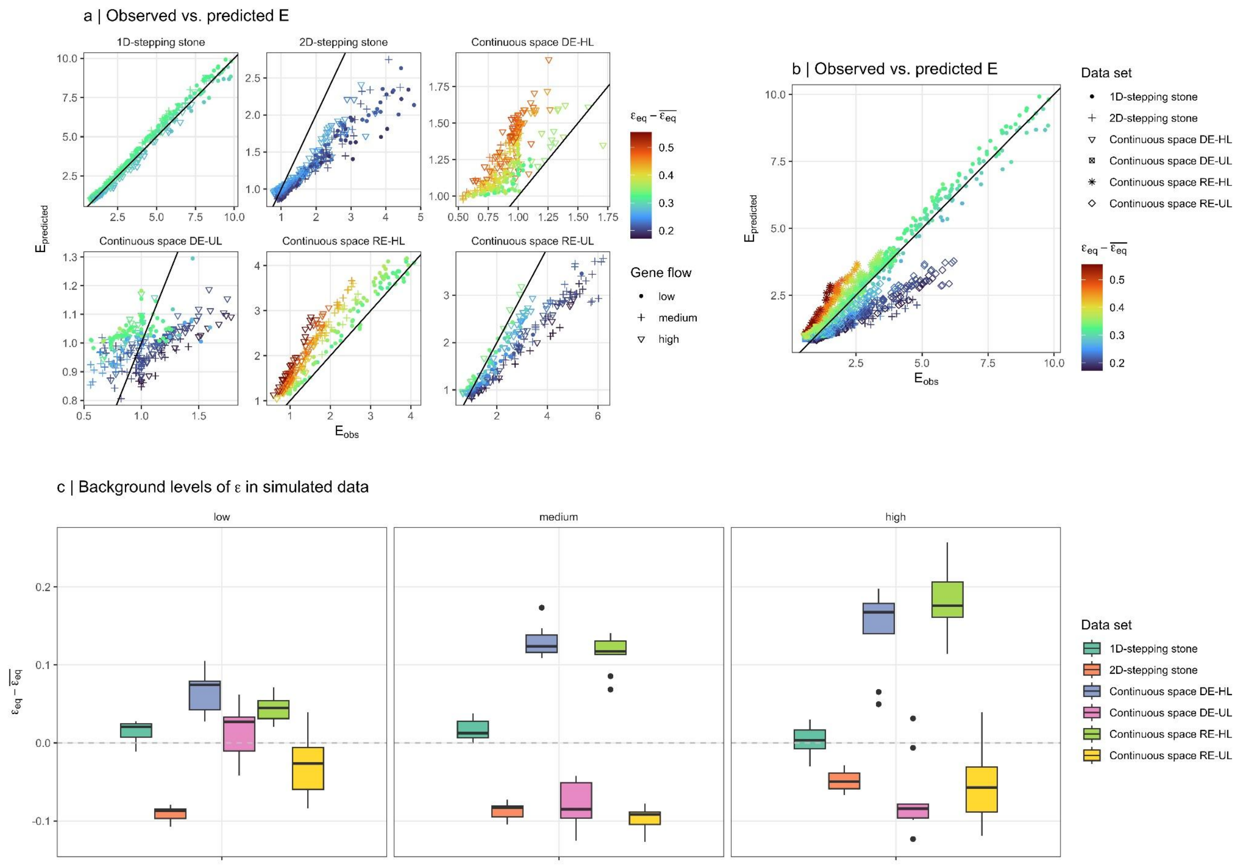


**Supplementary Figure S9 | Background levels of ϵ in simulated data.** Figure (a) and (b) shows how well the full model (2-main document) predicts *E* in different simulation data sets. In (a) x- and y-scales are differ between the panels and in (b) all values are shown in the same graph. The color indicates $\epsilon_{eq} - \overline{\epsilon_{eq}}$, where $\epsilon_{eq}$ is the equilibrium value of $\epsilon = |\psi|/\overline{F}_{ST}$ in each individual data set measured at the end of the simulations and $\overline{\epsilon_{eq}}$, is the mean across all data sets. In (b) the mean $\epsilon_{eq}$ is shown for all simulations, showing that the background level of ϵ (due to boundary effects) varies more between simulation datasets than between different levels of gene flow. When $\epsilon_{eq} - \overline{\epsilon_{eq}}$ is negative *E* will be underestimated relative to the true value of *E*. Consequently, among the simulated data sets, the power to detect *E*>1 is lower for data from the 2D-stepping stone and the continuous space simulations where the landscape is uniform (RE-UL; a,b,c).

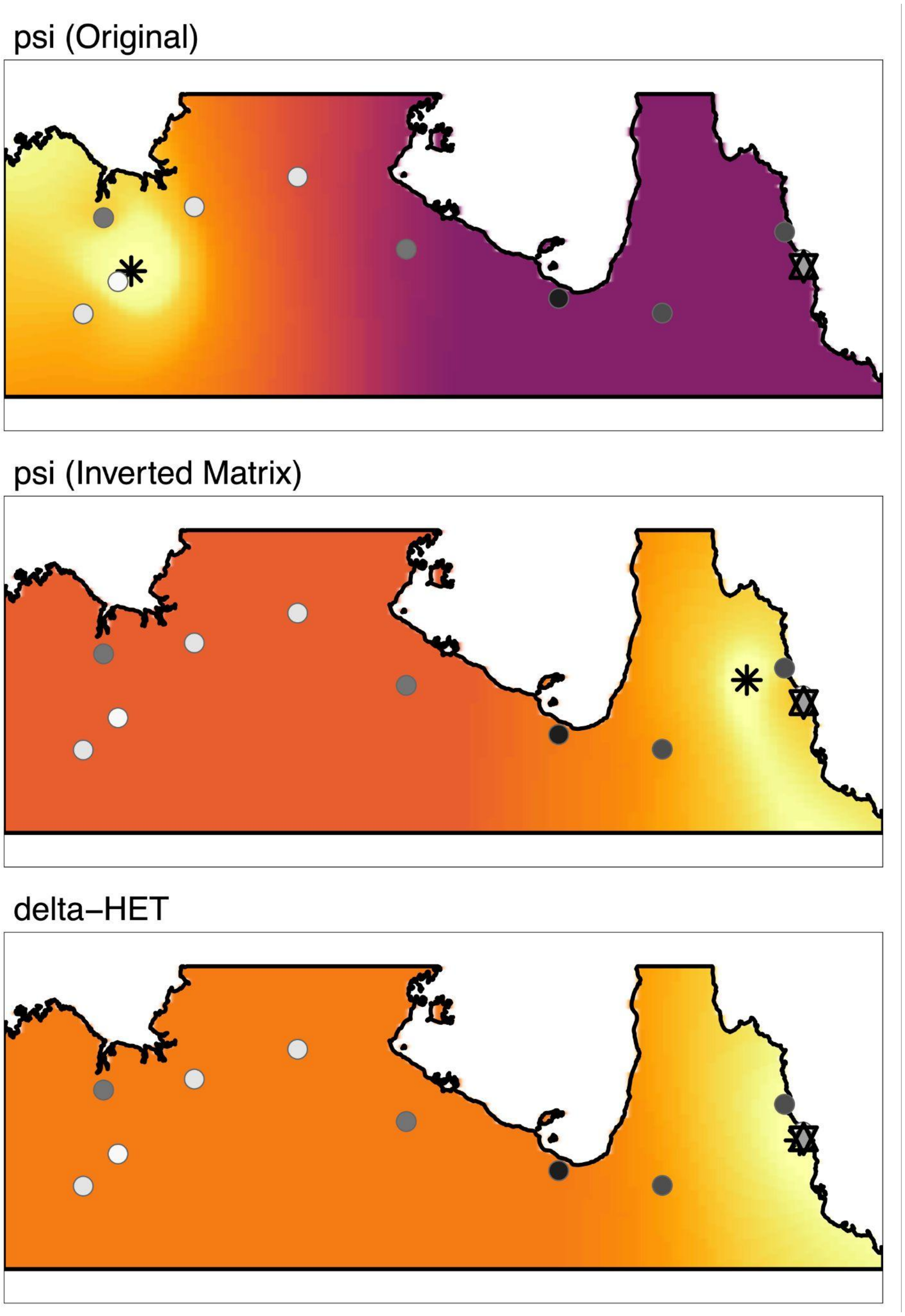


**Supplementary Figure S10 | Re-analyses of population genomic data from the invasion of cane toads in Northern Australia.** When using the original implementation of the function *get.all.psi* from the rangeExpansion R-package (top) the estimated origin is opposite to when the modified function *get.all.psi.mc.bin* was used. The latter resulted in the estimated origins (indicated by "*") being highly congruent with the known introduction site at Gordonvale, North Queensland (indicated by " ✡ ") using both ψ (middle) and $\Delta_{het}$ (bottom).
